# Metastatic organotropism in small cell lung cancer

**DOI:** 10.1101/2024.10.07.617066

**Authors:** Manan Krishnamurthy, Anjali Dhall, Sarthak Sahoo, Christopher W. Schultz, Michelle A. Baird, Parth Desai, Jacob Odell, Nobuyuki Takahashi, Michael Nirula, Sophie Zhuang, Yue Huang, Brett Schroeder, Yang Zhang, Maria Sebastian Thomas, Christophe Redon, Christina Robinson, Lai Thang, Lilia Ileva, Nimit L. Patel, Joseph D. Kalen, Alice-Anaïs Varlet, Noam Zuela-Sopilniak, Ankita Jha, Darawalee Wangsa, Donna Butcher, Tamara Morgan, Alyah N. Afzal, Raj Chari, Karim Baktiar, Suresh Kumar, Lorinc Pongor, Simone Difilippantonio, Mirit I. Aladjem, Yves Pommier, Mohit Kumar Jolly, Jan Lammerding, Ajit Kumar Sharma, Anish Thomas

## Abstract

Metastasis is the leading cause of cancer-related deaths, yet its regulatory mechanisms are not fully understood. Small-cell lung cancer (SCLC) is the most metastatic form of lung cancer, with most patients presenting with widespread disease, making it an ideal model for studying metastasis. However, the lack of suitable preclinical models has limited such studies. We utilized rapid autopsy-derived tumors to develop xenograft models that mimic key features of SCLC, including histopathology, rapid and widespread development of metastasis to the liver, brain, adrenal, bone marrow, and kidneys within weeks, and response to chemotherapy. By integrating in vivo lineage selection with comprehensive bulk and single cell multiomic profiling of transcriptomes and chromatin accessibility, we identified critical cellular programs driving metastatic organotropism to the liver and brain, the most common sites of SCLC metastasis. Our findings reveal the key role of nuclear-cytoskeletal interactions in SCLC liver metastasis. Specifically, the loss of the nuclear envelope protein lamin A/C, encoded by the *LMNA* gene, increased nuclear deformability and significantly increased the incidence of liver metastasis. Human liver metastases exhibited reduced *LMNA* expression compared to other metastatic sites, correlating with poorer patient outcomes and increased mortality. This study introduces novel preclinical models for SCLC metastasis and highlights pathways critical for organ-specific metastasis, offering new avenues for the development of targeted therapies to prevent or treat metastatic disease.

## Introduction

Metastasis is the leading cause of cancer-related mortality, but its biological underpinnings are still largely unknown. Understanding the molecular mechanisms that enable cancer cells to metastasize, i.e., ‘metastatic potential’ could inform strategies to prevent and treat this lethal phenomenon.

The metastatic cascade is driven by both tumor-cell intrinsic features and extrinsic cues from the microenvironment [1]. Tumor cells, enabled by genomic alterations that provide fitness advantages, metastasize via circulation after disseminating from primary sites. Pan-cancer studies have revealed a higher frequency of recurrent alterations in cancer driver genes, including *PIK3CA*, *APC*, *PTEN*, and *MYC*, and higher overall burden of copy number alterations, including whole genome duplication in metastases compared with primary tumors [2, 3]. Colonization of metastatic sites is also dependent on the compatibility of tumor cells with the local organ microenvironment, leading to the apparent organ preference or tropism of metastasis [4]. Tumor clones characterized by distinct transcriptional drivers disseminate at different rates to specific sites [5]. Therefore, knowledge not only of which cells metastasize, but also of organ-specific selective pressures that affect propensities to colonize different organs are important to reveal the full range of biological features that underpin metastasis.

Small-cell lung cancer (SCLC), a high-grade neuroendocrine cancer, is the most metastatic and fatal form of lung cancer. It is also arguably the most metastatic solid tumor, serving as a useful model for studying metastasis. Most SCLC patients present with widely metastatic disease, even when diagnosed using modern screening modalities [6]. SCLC primarily metastasizes to the liver, brain, bone, and adrenal glands, but the mechanisms driving its varied organotropism remain poorly understood [7] due to limited availability of metastatic tissue for research. SCLC is typically diagnosed using fine needle aspirates, and biopsies of metastatic tumors are not standard. Moreover, SCLC is absent from large-scale sequencing projects like The Cancer Genome Atlas. Clinically relevant models of SCLC metastasis are scarce. Despite the development of over 100 SCLC cell lines, only 7% (8/118) are derived from visceral metastatic sites [8]. Additionally, SCLC is not included among the 90+ brain-metastatic cancer models in the BrMPanel [9]. Further, it is unclear whether the few published human cell lines that metastasize upon mouse implantation retain relevant metastatic features from their cells of origin [7]. While genetically engineered mouse models of SCLC exist [10–12], their genetics are not fully representative of human disease, these tumors exhibit far lower metastatic propensity than human SCLC, and none reliably metastasize to the brain [7, 13, 14].

Research autopsies provide a unique opportunity for collection of a diverse set of samples that represent the full metastatic burden at the time of death. In this study, we introduce novel SCLC xenograft models developed from rapid autopsy-derived tumors. These models closely mirror the metastatic burden of the patient and, for the first time, captures organotropism, characteristic of SCLC metastasis. Using this model, we identify key transcriptomic and epigenomic drivers of metastasis, revealing the role of lamin A/C, encoded by the *LMNA* gene, highlighting its central role in promoting liver metastasis.

## Results

### Establishment and characterization of a patient-derived model of SCLC metastasis and organotropism

To address the challenge of limited SCLC metastasis models [7], we established a model using tumors from various metastatic sites obtained immediately after death from a 65 year old male patient (RA22) with advanced, treatment refractory SCLC. Tumor samples were obtained and cell lines generated from the lung (named 12-lung), cervical lymph node (4-lymph node), two distinct liver tumors (5-liver and 6-liver), and adrenal (18-adrenal) (**Fig. 1A**, Fig. S1A, Table S1) [15]. Cell lines generated from these tumors were directly injected into the left ventricle of NOD scid gamma (NSG) mice. Organ invasion and colonization was monitored weekly using magnetic resonance imaging (MRI) post-injection.

**Fig. 1:**
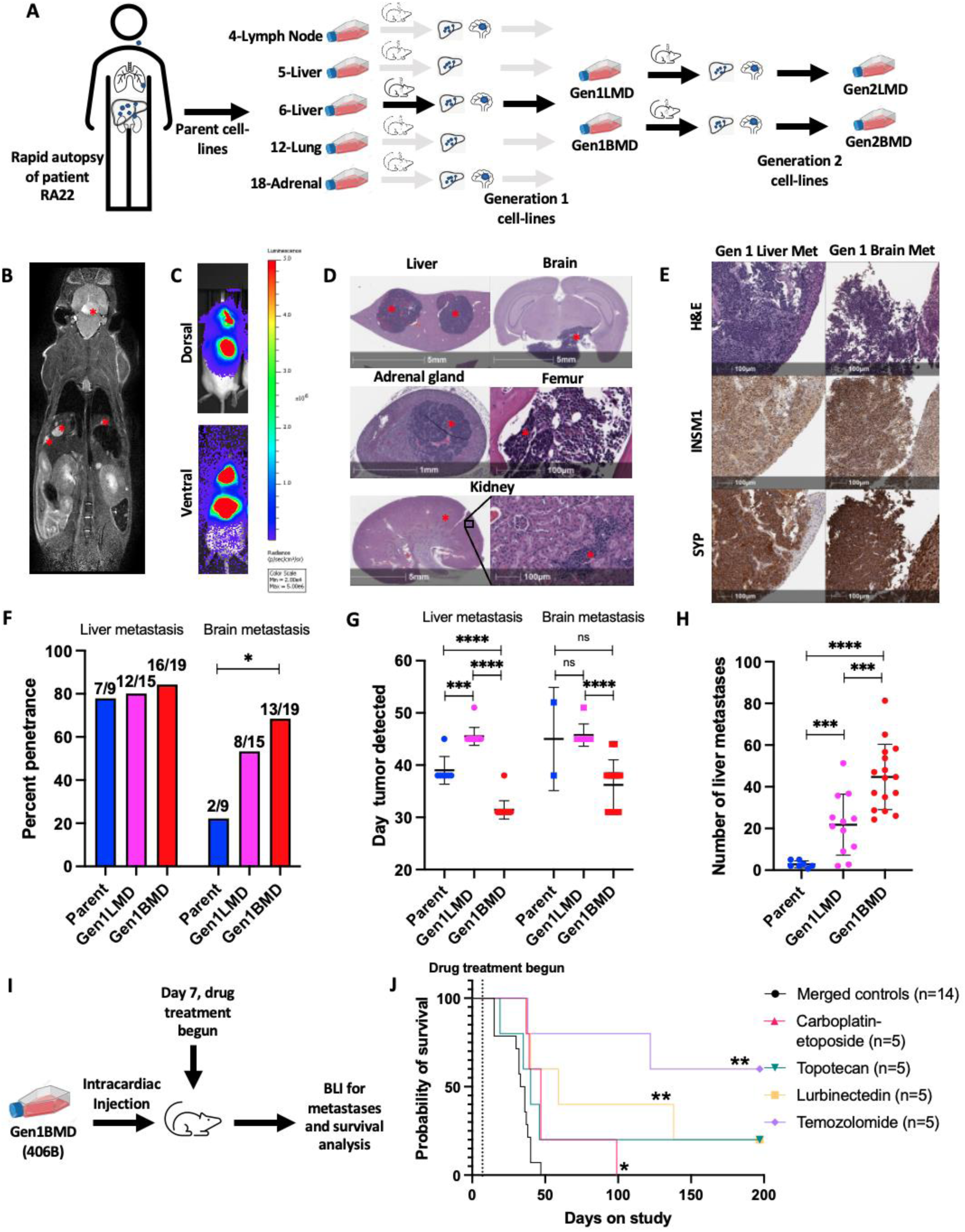
Establishment and characterization of a patient-derived model of SCLC metastases and organotropism. A: Schematic representation of model derivation. B: MRI depicting liver and brain metastases (red asterisks) C: BLI showing brain and liver metastases. D: H&E staining illustrating liver, brain, adrenal, bone marrow, and kidney metastases (red asterisk) E: H&E and IHC staining for neuroendocrine markers F: Percentage penetrance (% of mice that formed tumors) of brain and liver metastases across parental and in vivo selected liver and brain-tropic cell lines, assessed by MRI. Mice numbers indicated above each bar. Statistics by Fisher’s exact test. G: Time to develop liver and brain metastasis in days, assessed by MRI. Error bars: mean and standard deviation. H: Number of liver metastases, assessed by MRI. Error bars: mean and standard deviation. I: Schematic representation of drug assessment experiment J: Survival post intracardiac cell injection with and without treatment with standard SCLC therapeutics, Gen1BMD cell line 406B, n=5 per treatment group, n=14 merged controls (see Fig. S1R (merged control) and Table 1 for details). Statistics by Mantel-Cox test. X axes of figures F-H refer to the cell line injected. Significance tests used parametric unpaired T-tests with Welch’s correction and error bars are mean with standard deviation unless mentioned otherwise. *p<0.05, **p<0.01, ***p<0.001, ****p<0.0001. Abbreviations: LMD: Liver metastasis derived; BMD: Brain metastasis derived; BLI: bioluminescent Imaging; MRI: magnetic resonance imaging; SCLC: small cell lung cancer; Gen1LMD includes cell lines 400L and 404L; Gen1BMD includes cell lines 406B and 408B; Gen2LMD includes cell lines 431L and 431L; Gen2BMD includes cell lines 438L and 438B.

**Table 1:**

| Treatment | # of mice | Vehicle | Route | Dosing schedule |
| --- | --- | --- | --- | --- |
| Vehicle Control (Topotecan) | 7 | Saline | IP | M-F, one week off then another M-F |
| Vehicle Control (TMZ) | 7 | 20% hydroxypropyl-β-cyclodextrin | PO 2x day | M-F for 3 weeks |
| Vehicle Control (Lurbinectedin) | 7 | Saline | IV | Day 1, one week off then day 15 |
| 50 mg/kg Carboplatin | 5 | Saline | IP | Day 1, one week off then day 15 |
| 8 mg/kg Etoposide |  | Saline | IP | Days 1, 2, and 3 one week off then days 15, 16, 17 |
| 1 mg/kg Topotecan | 5 | Saline | IP | Week 1 M-F, one week off then Week 3 M-F |
| 0.18mg/kg Lurbinectedin | 5 | Saline | IV | Day 1, one week off then day 15 |
| 50 mg/kg Temozolomide | 5 | 10% DMSO, 90% D5W | PO | M-F, one week off then another M-F |

We observed metastatic tumors in liver and brain, mirroring the sites of metastases observed in the patient (**Fig. 1B**, Fig. S1B-G). Liver metastases were numerous and distributed across all lobes (**Fig. 1B,C**, Fig. S1B,C). Brain metastasis typically manifested as solitary lesions that invaded the pituitary, effacing, replacing, and expanding the architecture, extending into the meninges and invading the neuropil parenchyma (**Fig. 1D**, Fig. S1F,G). In some instances, brain metastases were located higher amidst the midbrain, cerebellum, and pons (Fig. S1F). A detailed examination of tissue sections using hematoxylin and eosin (H&E) staining revealed additional metastases in the kidneys, adrenal, and the marrow of the femur (**Fig. 1D**). Adrenal metastases were often indistinguishable from normal adrenal tissue on MRI (Fig. S1H). Histological analysis showed that all metastases consisted of tumor cells with small size, scant cytoplasm, and granular nuclear chromatin, characteristic of SCLC (**Fig. 1D, E**, Fig SG). Immunohistochemical staining confirmed positivity for multiple diagnostic SCLC markers, including insulinoma1 (INSM1), synaptophysin (SYP), chromogranin A, CD56, and Ki-67 (**Fig 1E**, Table S2, Fig. S1I).

Despite originating from the same patient, significant variation was observed in terms of tropism to the liver and brain (Fig. S1J). While all parent cell lines successfully colonized the liver, penetrance, the percentage of injected mice that developed liver metastases, varied widely: 80% for lymph node, 70% and 78% for liver, 75% for lung, and 22% for adrenal-derived cell lines (Fig S1J). On average, liver metastases became detectable on MRI between 38 to 40 days after injection of 4-lymph node, 6-liver, and 12-lung, whereas 5-liver and 18-adrenal exhibited metastases with an average latency of 45 to 49 days (Fig. S1K). 4-Lymph node and 12-lung demonstrated the highest liver tumor burden, averaging 25 to 35 tumors per mouse, while 5-liver and 18-adrenal formed fewer liver tumors, with approximately 3 to 4 tumors per mouse (Fig. S1L). 4-Lymph node and 12-lung exhibited rapid metastasis with a large total number of lesions, whereas 6-liver metastasized quickly but in smaller numbers.

Variation in brain tropism was evident across cell lines from different metastatic sites. Brain metastases occurred in 4-lymph node, 18-adrenal, and 6-liver, but not in 5-liver or 12-lung (Fig. S1J). Penetrance for brain metastasis was lower than liver metastasis, ranging from 50% for 4-lymph node to 22% for 5-liver, 6-liver, and 18-adrenal. Latencies varied, with 4-lymph node injection yielding brain metastases between days 38 and 45, 6-liver between days 38 and 52, and 18-adrenal on day 45 (Fig. S1K). Notably, brain metastases appeared within weeks, reflecting SCLC’s strong propensity for brain colonization, unlike models of other tumor types which require multiple rounds of mouse injections [16–21].

The observation that 4-lymph node exhibited the highest metastatic potential to both the brain and liver aligns with the well-known propensity of lymph node metastases to serve as a source of distant metastases [22]. Conversely, the cell line derived from adrenal displayed the longest overall latency, low penetrance, and low liver metastatic burden. For further investigations, we selected the 6-liver (henceforth called Parent), due to its metastasis to both the liver and brain with short latency, and high penetrance to the liver (Fig. S1J-L). In summary, our patient-derived model successfully recapitulated key features of SCLC, including organotropism, histomorphology, numerous macrometastases to the liver and brain, less common macrometastases to the adrenal and bone marrow, and micrometastases to the kidney.

### In vivo selection to enrich for tumor-cell populations with enhanced metastatic activity

To enrich for tumor cell characteristics associated with enhanced metastatic activity [23] (**Fig. 1A**, Fig. S1M), we performed intracardiac injection of Parent and derivative cell lines from brain and liver metastases. We repeated in vivo selection using two liver-derived and two brain-derived cell lines (named Gen1LMD and Gen1BMD respectively). Prior to reinjection, we tagged these lines with luciferase for downstream detection via bioluminescence imaging (BLI) (**Fig. 1A, C**). MRI was also performed to ensure that BLI detection correlated with macrometastases.

Gen1LMD and Gen1BMD demonstrated similar liver penetrance and improved brain penetrance as compared to Parent. Gen1BMD exhibited similar liver penetrance and improved brain penetrance compared to Gen1LMD (**Fig 1F**, Fig S1N). On average, Gen1BMD developed liver and brain metastases by day 31 and day 36, respectively, whereas Gen1LMD took until day 45 for both liver and brain metastases to develop (**Fig. 1G**, Fig. S1O). Gen1BMD also produced an average of 45 liver metastases, contrasting with 3 and 22 metastases each following injections of Parent and Gen1LMD (**Fig. 1H**, Fig. S1P). Overall, Gen1BMD exhibited the highest metastatic penetrance, shortest latency, and greatest metastatic burden, surpassing both Parent and Gen1LMD cells, thus emerging as the most rapid and penetrant model of SCLC metastasis. From the two Gen1BMD cells, we selected 406B, which displayed 100% metastatic penetrance to the liver and close to 90% penetrance to the brain for further selection and establishment of Gen2LMD and Gen2BMD variants. (Fig S1N).

Using the 406B model, we assessed the efficacy of standard of care SCLC therapeutics [24] including the carboplatin-etoposide combination typically used in newly diagnosed SCLC patients, as well as second-line treatments such as topotecan, lurbinectedin, and temozolomide (TMZ) to assess their ability to prevent metastatic colonization in vivo (**Fig1I**, Table 1). Drugs were administered 7 days after intracardiac injection of cells and survival following drug treatment was recorded until day 197. All treatment arms delayed both liver and brain metastatic progression and extended the average cohort survival compared to merged controls (**Fig. 1J**, Fig. S1Q-T). Remarkably, TMZ completely inhibited the formation of both liver and brain metastases with no tumor detected over the course of the experiment. Consistent with this high efficacy, expression of the *MGMT* (O6-methylguanine–DNA methyltransferase) DNA-repair protein, a major contributor to TMZ resistance [25] was undetectable in the patient tumor (Fig S1U).

In summary, we characterize a novel patient-derived model that faithfully recapitulates human SCLC metastases and demonstrates organ tropism. The model could serve as a tool to dissect the underlying factors and pathways contributing to the enhanced metastatic capability and organ specificity observed in SCLC.

### Transcriptomic alterations in SCLC liver and brain metastases

To explore the transcriptomic changes associated with SCLC metastatic organotropism, we performed bulk RNA-seq on Parent, Gen1LMD and Gen1BMD, as well as the Gen2LMD and Gen2BMD. Principal component analysis (PCA) of 2756 differentially expressed genes (DEG) revealed distinct separation of Gen1LMD and Gen1BMD from the Parent (**Fig. 2A**), indicating organ-specific evolution in liver and brain. PCA of DEGs showed co-clustering of Gen1BMD and its derivatives Gen2LMD and Gen2BMD away from Parent and Gen1LMD (**Fig. 2B**). This suggests that Gen2LMD retained transcriptomic features of brain metastases from which it was derived, rather than undergoing reprogramming to resemble the liver-adapted Gen1LMD. PCA of all genes illustrated similar trends with a stronger overlap between Gen1BMD, Gen2BMD, and Gen2LMD (Fig. S2A,B). Quantification of unique DEGs further revealed that Gen1LMD exhibited the highest number of unique transcriptomic changes, underscoring significant adaptations in the liver metastases (Fig. S2C,D).

**Figure 2:**
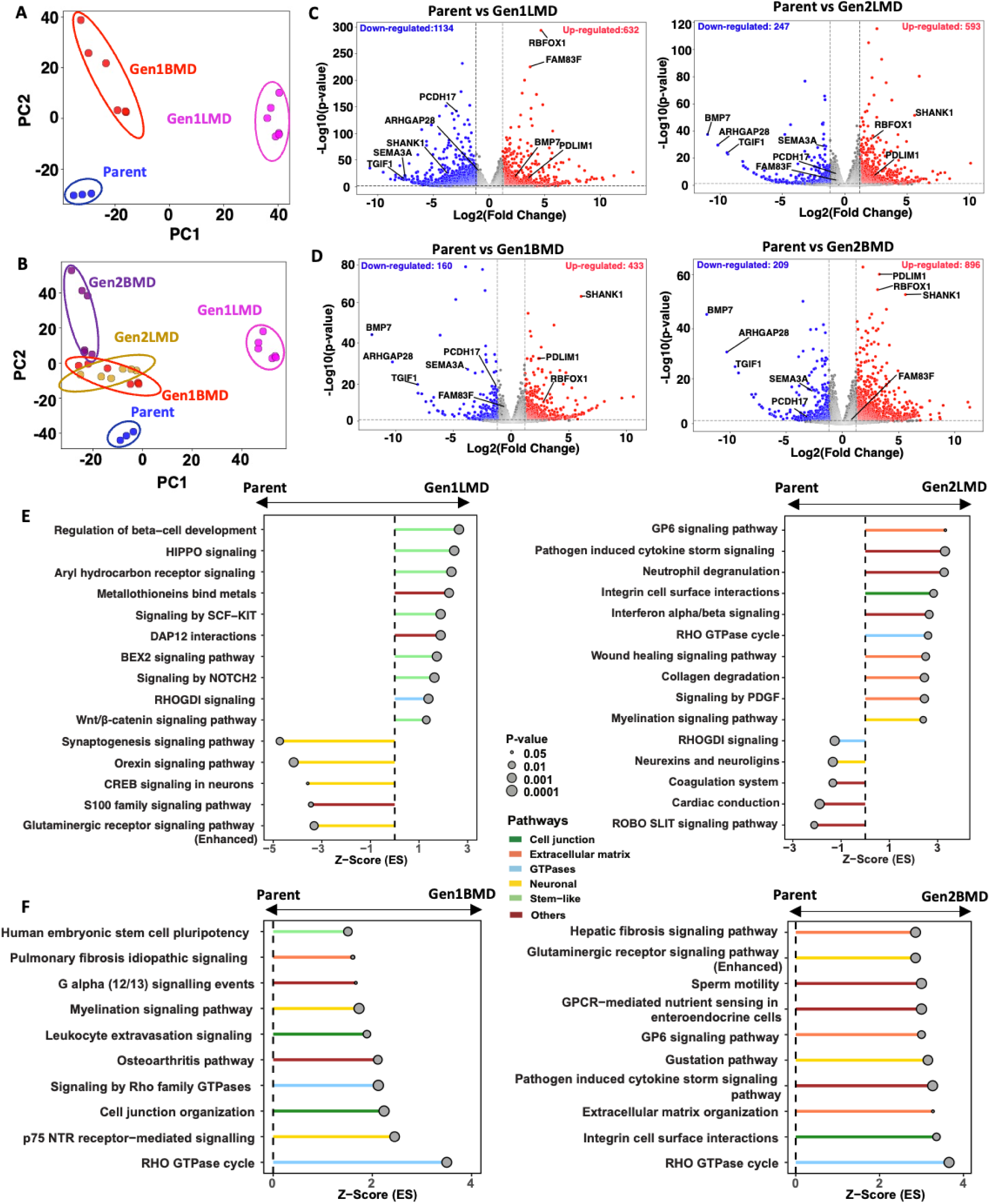
Transcriptomic alterations in SCLC liver and brain metastases. A: PCA of Parent and Gen1 differentially expressed genes, RNA-seq B: PCA of Parent, Gen1, and Gen2 differentially expressed genes, RNA-seq C: Pathways enriched in Gen1LMD (left) and Gen2LMD (right) compared to Parent D: Pathways enriched in Gen1BMD (left) and Gen2BMD (right) compared to Parent E: Volcano plots illustrating differentially expressed genes in Gen1LMD and Gen2LMD relative to Parent F: Volcano plots illustrating differentially expressed genes in Gen1BMD and Gen2BMD relative to Parent Abbreviations: PCA: Principal component analysis; IPA: Ingenuity pathway analysis; gen: generation; LMD: Liver metastasis derived; BMD: Brain metastasis derived

Differential gene expression analysis revealed key genes associated with liver and brain metastasis. Gen1LMD uniquely upregulated *FAM83F*, a WNT signaling regulator [26], and downregulated tumor suppressor *PCDH17*, which inhibits WNT signaling and metastasis [27] **(Fig. 2C**). Gen1BMD and Gen2BMD consistently upregulated *SHANK1*, a neuronal development protein, as compared to Parent [28]. Consistently downregulated genes in Gen1BMD and Gen2BMD included *BMP7*, an anti-metastatic factor involved in maintaining the epithelial phenotype [29], and *ARHGAP28*, whose deletion is associated with actin reorganization [30] (**Fig. 2D)**. Notably, DEGs in Gen2LMD largely mirrored Gen1BMD, suggesting conserved characteristics. All generations showed downregulation of tumor suppressor *TGIF1* [31] and cytoskeletal regulator *SEMA3A* [32–35]. All generations also showed upregulation of *RBFOX1* [36] and *PDLIM1*[37], involved in cytoskeletal regulation.

*MYC* RNA was also highly expressed in all generations (Fig. S2E), aligning with the significant levels of MYC amplification observed in patient tumors [15]. However, the patient tumor exhibited variability in the form of *MYC* amplification, with some instances detected as extrachromosomal DNA (ecDNA) and others as a homogeneously staining region (HSR) [15]. Further analysis revealed that *MYC* amplification was consistently maintained as an HSR across the Parent and its derivatives (Fig. S2F).

To identify the cellular pathways that characterize liver and brain metastasis, we employed Ingenuity Pathway Analysis (IPA) [38]. Relative to Parent, multiple pathways linked to stemness were significantly activated in Gen1LMD (**Fig. 2E**). Notably, WNT-beta catenin signaling, important for liver cell renewal and regeneration [39] and HIPPO signaling, essential for determining liver cell fate and maintaining plasticity [40] were both upregulated in Gen1LMD. RhoGDI signaling, important for spatial determination of the actin cytoskeleton was also upregulated [41]. In contrast, pathways related to neuronal signaling were downregulated in Gen1LMD. In Gen2LMD, the altered pathways differed substantially from those enriched in Gen1LMD (**Fig. 2E, F**), instead closely resembling those observed in Gen1BMD, the parent cell line of Gen2LMD (Fig. 1A), reflecting the distinct characteristics of metastatic generations and their lineage relationships.

In Gen1BMD and Gen2BMD, pathways significantly activated compared to the Parent were associated with neuronal signaling mechanisms (e.g. myelination), cell-cell junctions, ECM, stemness, and Rho GTPases (**Fig. 2F**). These findings are supported by recent reports of SCLC brain metastases co-opting mechanisms involved in neuron-astrocyte interactions [42]. Furthermore, disruption of cell adhesion and ECM degradation are critical processes in tumor colonization and adaptability at metastatic sites [43–45]. Remarkably, Gen2BMD displayed enhanced enrichment of ECM pathways compared to Gen1BMD, suggesting their increased importance in brain colonization. Lastly, Rho GTPase-associated pathways, canonically involved in cytoskeletal remodeling [46], cellular migration [47], and metastasis [48] showed significant alterations in BMD lines.

Overall, our transcriptomic data reveal genes and cellular programs linked to organ-specific metastatic behaviors at the most frequent sites of SCLC metastasis, liver and brain. The consistent expression of stem-like pathways in liver metastases, particularly the WNT-beta catenin signaling, along with pathways associated with cell-cell junctions and the ECM in brain metastases, implies distinct roles for these pathways in liver and brain-specific adaptations, respectively. Interestingly, cytoskeleton associated pathways and genes were differentially regulated in both liver and brain metastases. Importantly, this analysis suggests Gen1 cell lines may better recapitulate organ specific difference between brain and liver metastases as Gen2 lines appear to more resemble their Gen1 parent.

### Epigenomic alterations in SCLC organ-specific metastases

To evaluate the epigenomic changes associated with organ-specific metastases, we performed ATAC-seq on the Parent (6-Liver) and selected derivative cell lines: Gen1LMD (404L), Gen1BMD (406B), Gen2LMD (431L), and Gen2BMD (431B). PCA of differentially open peaks and all peaks relative to Parent (**Fig. 3A**, Fig. S3A) revealed distinct separation of Gen1LMD and Gen1BMD from Parent, mirroring findings from RNA-seq data (Fig. 2A, S2A).

**Figure 3:**
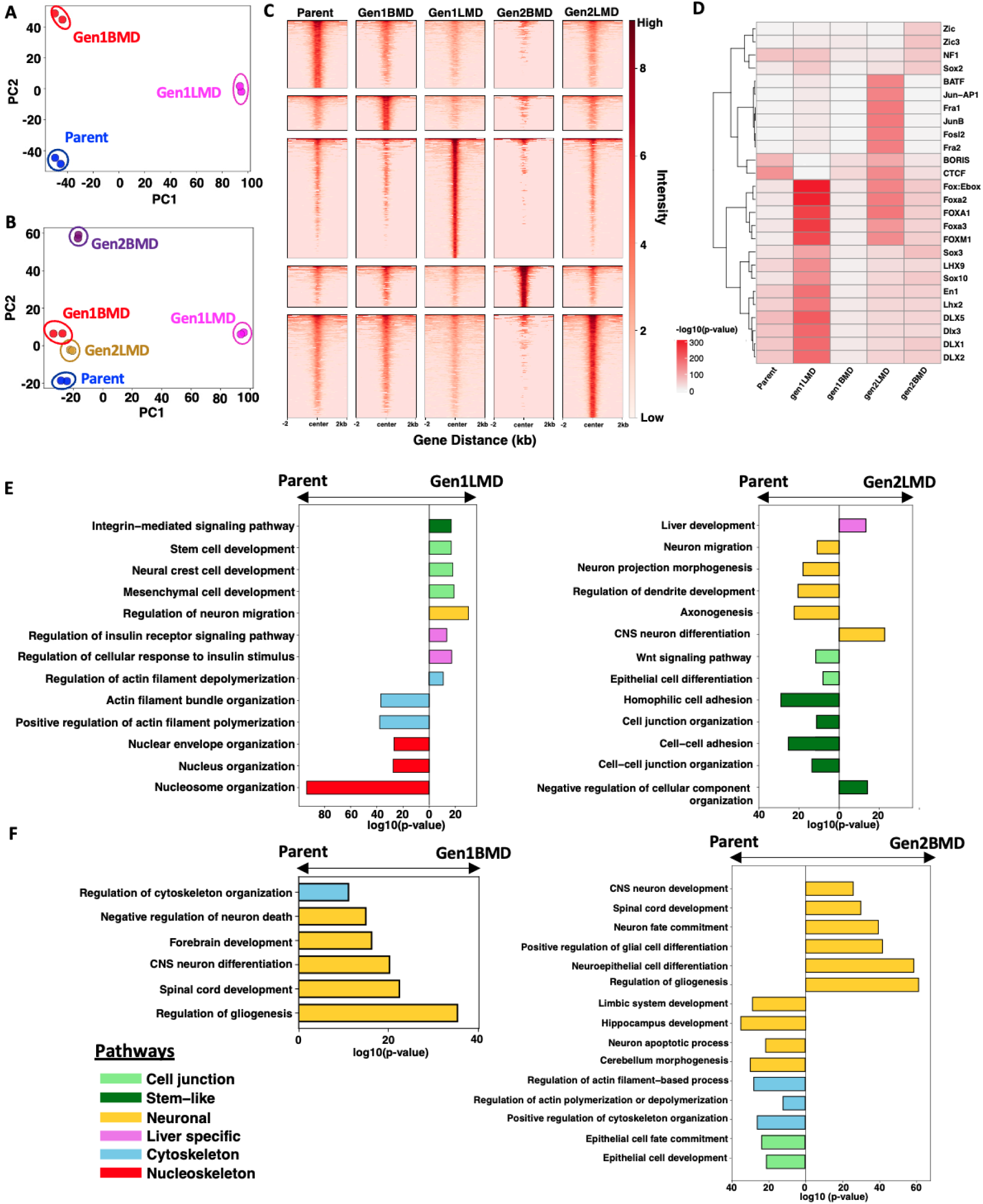
Epigenomic alterations in SCLC liver and brain metastases. A: Principal component analysis (PCA) of differential peaks from ATAC-seq in Parent and Gen1. B: PCA of differential peaks from ATAC-seq across Parent, Gen1, and Gen2. C: Heatmap of unique peaks in Parent, Gen1, and Gen2. D: Top 10 unique transcription factor motifs enriched in Parent, Gen1, and Gen2 E: Pathways potentially impacted by newly opened regions in Gen1LMD (left) and Gen2LMD (right) compared to Parent, GREAT analysis of differentially accessible peaks. n = number of observed peaks; arrows indicate which cell line had greater peak enrichment. F: Pathways potentially impacted by newly opened regions in Gen1BMD (left) and Gen2BMD (right) compared to Parent, GREAT analysis of differentially accessible peaks. n = number of observed peaks; arrows indicate which cell line had greater peak enrichment. Abbreviations: PCA: Principal component analysis; Gen, generation; LMD: Liver metastasis derived; BMD: Brain metastasis derived, GREAT, Genomic Regions Enrichment of Annotations Tool

A broader PCA that included Parent and all derivative cell lines similarly demonstrated clustering of Gen1BMD and its derivatives, Gen2LMD and Gen2BMD, away from Parent and Gen1LMD (**Fig. 3B**, 2B, S2B, S3B). Pairwise correlation analysis of chromatin accessibility further highlighted the distinct epigenomic profile of Gen1LMD, suggesting significant epigenomic reprogramming in the liver (Fig. S3C). Supporting this, Gen1LMD and Gen2LMD exhibited the highest number of unique open chromatin regions compared to both the Parent and the BMD-derived cell lines (**Fig. 3C**, S3D, F).

To identify key regulatory elements, we analyzed transcription factor motifs enriched in accessible chromatin regions. FoxA1/2 motifs were consistently enriched across all conditions, aligning with established SCLC oncogenic programs (**Fig. 3D**) [49]. Notably, FoxA1/2 enrichment was highest in Gen1LMD and Gen2LMD, potentially reflecting their roles specifically in liver metastasis development and adaptation [50].

We utilized the Genomic Regions Enrichment of Annotations Tool (GREAT) [51, 52] to examine pathways associated with the differentially open chromatin regions. This analysis revealed trends similar to those observed in the RNA-seq data, highlighting significant enrichment of stemness-related pathways in Gen1LMD and dysregulation of cytoskeletal reorganization signatures compared to the Parent (**Fig. 3E**). Additionally, regions linked to liver metabolic functions, particularly insulin response pathways, were enriched in Gen1LMD. Notably, the most significantly downregulated pathways in Gen1LMD were related to nuclear structure, suggesting that the disruption of genes maintaining nuclear integrity may facilitate metastasis [53]. In contrast, Gen2LMD exhibited de-enrichment of open peaks related to neuronal differentiation and cell junction pathways, with stem-like pathways, including Wnt signaling also de-enriched in Gen2LMD. Interestingly, a liver development pathway was enriched in Gen2LMD, indicating that adaptation to the liver microenvironment is more evident at the epigenomic level. Thus, while both Gen1LMD and Gen2LMD show enriched liver-specific characteristics, Gen2LMD adopts distinct characteristics likely influenced by its derivation from Gen1BMD.

In Gen1BMD, compared to the Parent, open peaks were enriched for neuronal functions and cytoskeletal organization regulation (**Fig. 3F**). In Gen2BMD, pathways involved in neuronal differentiation were further enriched, while pathways related to specialized brain regions and neuronal apoptotic processes were de-enriched. Despite the transcriptomic data suggesting extracellular matrix factor involvement, no corresponding enrichment was observed at the epigenomic level in any of the samples.

Overall, this analysis reveals epigenetic programs associated with organ-specific metastasis in the liver and brain. In brain, extensive enrichment of neuronal functions and variable accessibility of regions involved in cytoskeletal maintenance imply that distinct cellular mechanisms are crucial. Conversely, the enrichment of stem-like and metabolic pathways, coupled with de-enrichment of cytoskeletal and nuclear organization pathways suggests that these mechanisms may be vital for liver-specific metastasis.

### Single cell multiomic profiling of transcriptomes and chromatin accessibility in liver and brain metastases

To evaluate the transcriptomic and epigenomic heterogeneity associated with organ-specific metastases, we performed single-cell multiome RNA-ATAC sequencing on Parent (6-Liver) and selected derivative cell lines, Gen1LMD (404L) and Gen1BMD (406B). Two dimensional UMAP plots clearly separated Parent, Gen1LMD, and Gen1BMD (**Fig. 4A**), consistent with patterns observed in both bulk RNA and ATAC-seq analyses (**Fig. 2A, 3A**). Gen1BMD- and Gen1LMD-specific gene signatures derived from bulk data were appropriately enriched in the corresponding conditions in the single-cell data (**Fig. 4B**). As before, this analysis demonstrated that Parent shared greater similarity with Gen1BMD than with Gen1LMD.

**Fig 4.**
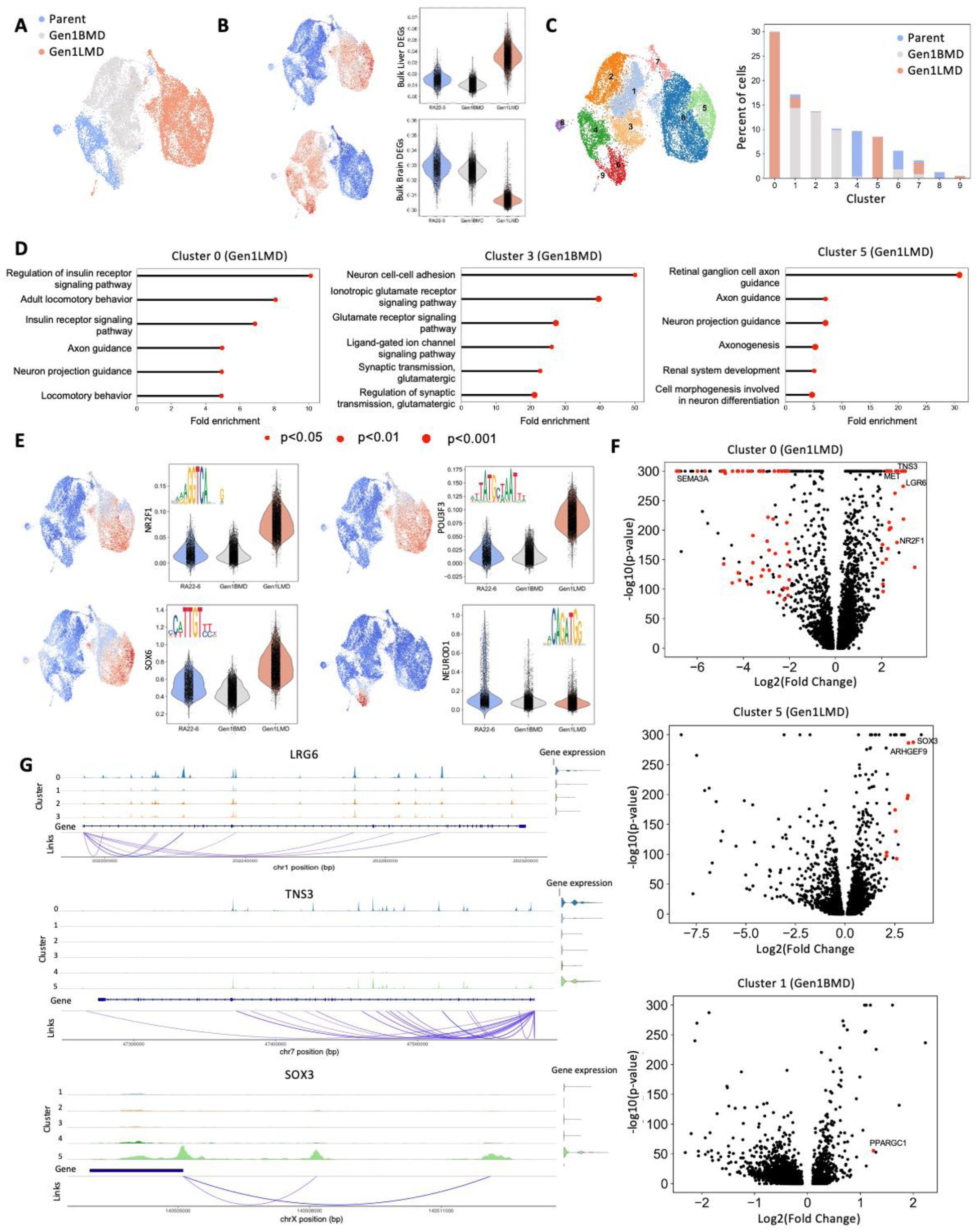
Single cell multiomic profiling of transcriptomes and chromatin accessibility in liver and brain metastases. A: UMAP of single cell multiome ATAC+ gene expression, Parent, Gen1LMD, and Gen1BMD. B: GSEA of Gen1LMD- and Gen1BMD-signatures from bulk RNA-seq. C: Cluster analysis and distribution of cells (shown as percent of all cells) within each cluster. D: Top gene ontology signatures of Gen1LMD (Cluster 0, 5) and Gen1BMD (Cluster 3). Gen1BMD Clusters 1 and 2 not yield significant pathway signatures; clusters 8 and 9 had few cells. Clusters 3, 6, and 7 in Fig S4A. E: Selected genes that showed differentially open peaks and elevated expression in multiple Gen1LMD clusters and in Cluster 6 that was shared between Parent and Gen1BMD cells. Gen1BMD did not have any factors that fit these criteria. F: Volcano plots of differentially expressed genes in Gen1LMD and Gen1BMD. Red dots indicate genes that also show epigenetic linkage. y cut off at -log10(p-value) = 300 and all remaining genes represented in one line. G: Representative coverage plots for LRG6, TNS3, and SOX3. Full coverage plots for all called out genes in Fig. S4

Next, cluster analysis revealed distinct cell states among Parent (clusters 4, 6, 8, 9), Gen1LMD (clusters 0 and 5), and Gen1BMD (clusters 1, 2, 3) (**Fig. 4C**). Cluster 6 was shared between Parent and Gen1BMD cells, while Cluster 7 comprised a mix of all conditions. Gen1LMD-specific clusters 0 and 5 were enriched for neuronal pathway signatures, consistent with observations in SCLC metastases (**Fig. 4D**) [54, 55]. Notably, Cluster 0 also exhibited an insulin regulation signature, suggesting additional adaptation to the liver niche exists but is not essential for all cell states. Gen1BMD-specific Cluster 3 was enriched for neuron adhesion and neurotransmitter synapse signatures. Shared Cluster 7 cells demonstrated enrichment in cell-cycle pathways (Fig. S4A). Clusters 1 and 2 did not yield sufficient DEGs for analysis, and clusters 8 and 9 had too few cells for meaningful interpretation.

To further explore epigenomic regulation across the clusters, we identified motifs that were enriched in each cluster (Fig. S4B). We then examined whether transcription factors binding these motifs exhibited increased gene expression across multiple clusters in each condition. Notably, liver-derived Gen1LMD cells showed differentially open peaks and elevated expression of *NR2F1*, *POU3F3*, and *SOX6*, transcription factors that are critical for neural cell development [56–59] (**Fig. 4E**). This finding highlights the role of neuronal stemness in liver metastasis of SCLC. In contrast, no such transcription factors were identified in the Gen1BMD-specific clusters, potentially due to the lower coverage inherent to single-cell sequencing. Interestingly, Cluster 6 which was shared between Parent and Gen1BMD cells showed differentially open peaks and elevated expression of *NEUROD1*.

Next, we identified DEGs within each cluster and selected genes with open chromatin regions corresponding to their expression levels (indicated by red dots) (**Fig. 4F**, Fig. S4C, Table S3). This approach enabled the identification of accessible genomic regions potentially involved in gene regulation.

In Gen1LMD cluster 0, upregulated genes included *LRG6*, *MET* and *TNS3*, while *SEMA3A* was downregulated (**Fig. 4F**). The stem cell marker LRG6 has been shown to promote metastasis via enhanced WNT-beta catenin signaling, a pathway also observed in Gen1LMD bulk RNA-seq (**Fig. 2E**, **Fig. 4G**) [60]. *MET* binds hepatocyte growth factor and plays a role in cell motility, stem-like phenotypes, tumor progression, and dysregulation of cytoskeletal regulation of nuclear shape [61–63]. *TNS3* supports cell migration and cancer metastasis [64]. In contrast, cytoskeletal regulator *SEMA3A* downregulation is associated with enhanced tumor angiogenesis and progression in solid tumors [32–35].

In Gen1LMD cluster 5 *SOX3* and *ARHGEF9* were notably upregulated, whereas no linked genes were downregulated. SOX3 is associated with tumor progression in liver cancer and ARHGEF9 promotes cytoskeleton variations that determine cell share in melanoma (**Fig. 4F)** [65, 66]. Notably, in both Gen1LMD clusters 0 and 5 genes important for nucleo-cytoskeletal organization were dysregulated. Consistently, GSEA using bulk-ATAC data identified nuclear envelope organization signatures with decreased expression (Fig. S4D).

In contrast, Gen1BMD clusters exhibited fewer DEGs (**Fig. 4F**, Fig. S4C). In Gen1BMD Cluster 1, *PPARGC1A* was upregulated, a gene linked to fatty acid synthesis and metastatic outgrowth of brain metastases (Fig. S4G) [67]. Coverage plots illustrating epigenetic regulation and gene expression for the mentioned genes are shown (**Fig. 4G**, S4E-K). No epigenetic links were identified for clusters without corresponding volcano plots.

Overall, this analysis reveals the diverse epigenomic and transcriptomic adaptations underlying liver and brain metastases in our model. Liver metastases were characterized by two distinct epigenetically regulated cell clusters, each exhibiting unique transcriptional signatures but sharing common dysregulation in factors governing nucleo-cytoskeletal organization. In contrast, brain metastases were defined by transcriptional programs enriched for synaptic neuronal functions, reflecting a specialization for the neural microenvironment. Collectively, these results underscore the organ-specific epigenomic and transcriptional plasticity employed by SCLC metastases.

### Lamin A/C knockout enhances SCLC liver metastatic burden

Our finding of transcriptional and epigenetic dysregulation in cytoskeletal organization are consistent with the reduced cytoplasmic volume and distinct nuclear morphology of SCLC cells, known as nuclear molding – a diagnostic hallmark on histology (Fig. S5A) [70]. We identified dysregulation of several genes encoding cytoskeletal components, including *ACTB* (beta-actin) and *TUBB* (beta-tubulin), as well as key elements of the LINC (linker of nucleoskeleton and cytoskeleton) complex, *SUN1* and *SUN2*, which connect the cytoskeleton to the nuclear envelope (Fig. S5B). Additionally, *LMNA* (Lamin A/C), an essential nuclear envelope protein required for nuclear stability, nuclear-cytoskeletal coupling (in concert with the LINC complex), chromatin organization, and transcriptional regulation [71], was also dysregulated. In Gen1LMD, we observed significant downregulation of *LMNA* compared to the parental line (**Fig. 5A**), accompanied by reduced chromatin accessibility at the *LMNA* locus (**Fig. 5B**). In contrast, *LMNA* was upregulated in all brain metastasis-derived lines, suggesting its potentially metastatic site-dependent role in cancer progression [72].

**Fig 5:**
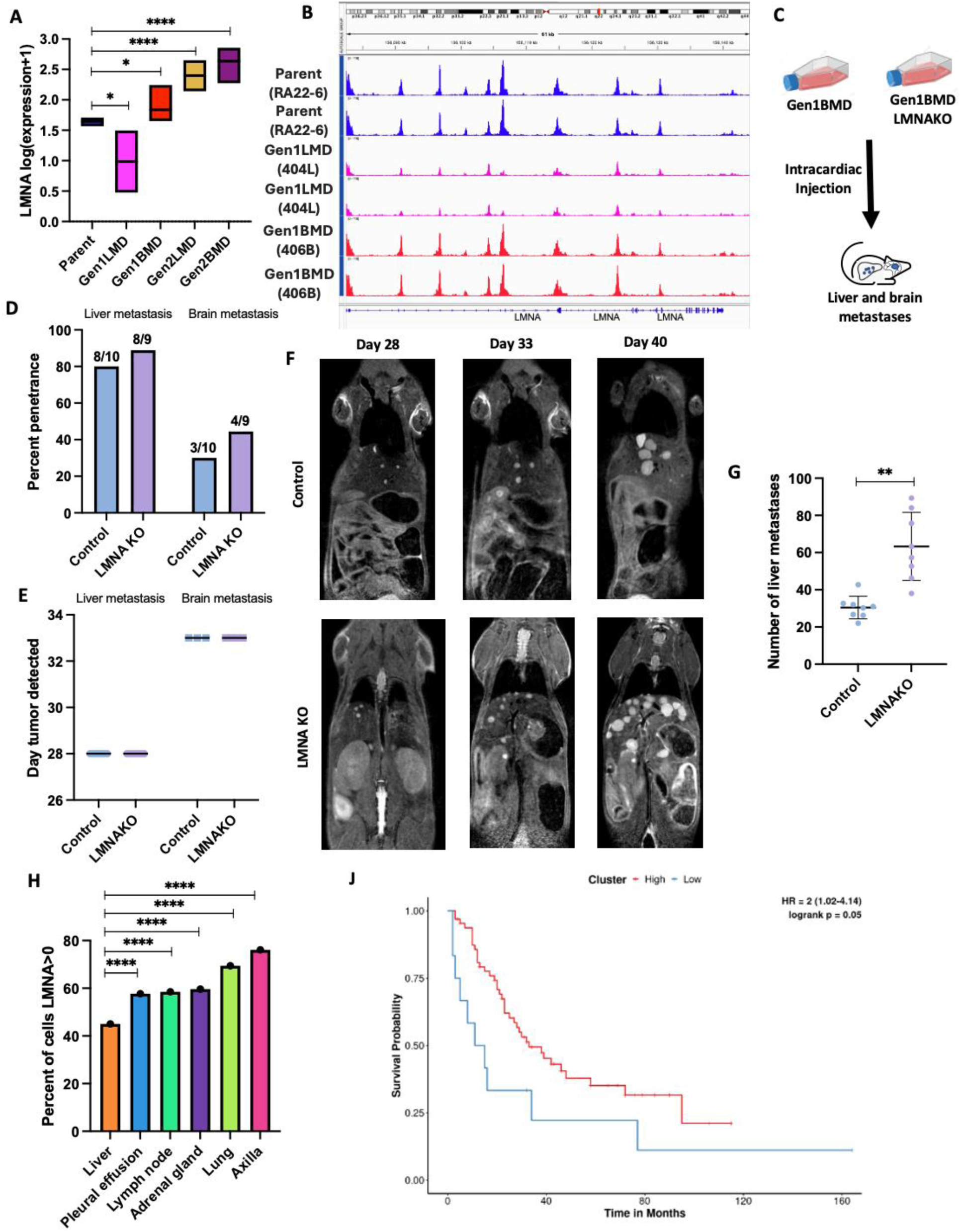
Lamin A knockout enhances SCLC liver metastatic burden. A: RNA-seq of LMNA gene in Parent, generation 1, and generation 2. t-tests only shown compared to Parent. B: ATAC-seq of LMNA locus in Parent and generation 1, two replicates each sample C: Schematic of mouse experiment D: Percentage penetrance (% of mice that formed tumors) of brain and liver metastases in 406B and 406BLMNAKO cell lines, assessed by MRI. Mice numbers indicated above each bar. E: Time to develop liver and brain metastasis in days, assessed by MRI. Error bars: mean and standard deviation. F: Representative images of liver metastases in control and LMNAKO, assessed by MRI. G: Quantification of number of liver metastases on day 33 assessed by MRI. Error bars: mean and standard deviation H: Percent of the 54,523 human SCLC cells [68] expressing LMNA, stratified by site of metastasis. Chi-squared tests only shown compared to liver. J: LMNA expression and survival (n=77 patients) [69], stratified lower 15th percentile of LMNA expression compared to the rest. Significance tests used parametric unpaired T-tests with Welch’s correction and error bars are mean with standard deviation unless mentioned otherwise. *p<0.05, **p<0.01, ****p<0.0001.

To further understand the role of LMNA in liver metastasis, we generated a CRISPR knockout of *LMNA* in our highly metastatic 406B SCLC cell line (LMNAKO) (Fig. S5C, D). RNA-seq of LMNAKO confirmed widespread transcriptional changes in LINC complex and cytoskeletal components, indicating alterations in nuclear mechanics (Fig. S5E). To evaluate whether LMNA loss enhanced metastatic potential, LMNAKO and control cells were injected intracardially into NSG mice (n=10 per group) (**Fig. 5C**). Both groups showed similar liver and brain metastasis penetrance (**Fig. 5D**) and similar latency periods (**Fig. 5E**). However, LMNAKO mice developed significantly more liver metastases (**Fig. 5F, G**), suggesting that LMNA loss enhances metastatic burden by promoting increased cell extravasation to or colonization of the liver, though it does not affect metastasis initiation. LMNA absence was verified in cell lines derived from metastases (Fig. S5F). Furthermore, the de-enrichment of nuclear envelope genes in Gen1LMD specifically aligns with the observed increase in liver metastasis, reinforcing the role of nuclear envelope dysregulation in promoting liver-specific colonization (**Fig. 3D**).

These data indicate that LMNA loss specifically promotes liver metastasis by enhancing extravasation or colonization at metastatic sites. To explore this further, we analyzed a single-cell RNA sequencing (scRNA-seq) dataset of 54,523 SCLC epithelial cells [68]. Only 45% of liver-derived cells expressed *LMNA*, compared to 69% of cells from primary lung tumors (**Fig. 5H**), and *LMNA*-expressing cells in the liver exhibited lower average expression levels than those from other metastatic sites (Fig. S5G). Supporting the correlation between reduced *LMNA* expression and increased liver metastasis potential, patients with the lowest 15th percentile of *LMNA* expression had significantly worse survival [69] (**Fig. 5J**).

### Lamin A knockout enhances SCLC deformability and migration through confined spaces

LMNA is a major determinant of nuclear deformability [73–75], which is considered a rate-limiting step in the migration of cancer cells through confined spaces during metastasis [76, 77]. Building on our observation that the loss of the nuclear envelope protein LMNA enhances liver metastatic burden, we aimed to quantify and compare nuclear deformability [78, 79] and the ability to migrate through confined spaces [80] between 406B LMNAKO and control cells. Assays using RA22 patient-derived cells were complicated by their clumped growth patterns, which interfered with adhesion assays and reliable measurements. To address this, we utilized the adherent human SCLC cell line DMS114 and generated an LMNAKO variant (Fig. S6A).

To investigate whether LMNA loss affects metastasis by enhancing nuclear deformability, we conducted time-lapse microscopy alongside a polydimethylsiloxane (PDMS) confinement assay (**Fig. 6A**) [78], which constrains cells to a height of 3 µm, simulating the confined ECM environments encountered during metastasis [77]. Hoechst-stained DMS114WT and LMNAKO cells were assessed for changes in nuclear perimeter pre- and post-confinement. The LMNAKO cells showed a more significant increase in nuclear perimeter (average 1.54-fold) than controls (average 1.30-fold) (**Fig. 6B, C**), indicating greater nuclear deformability.

**Figure 6:**
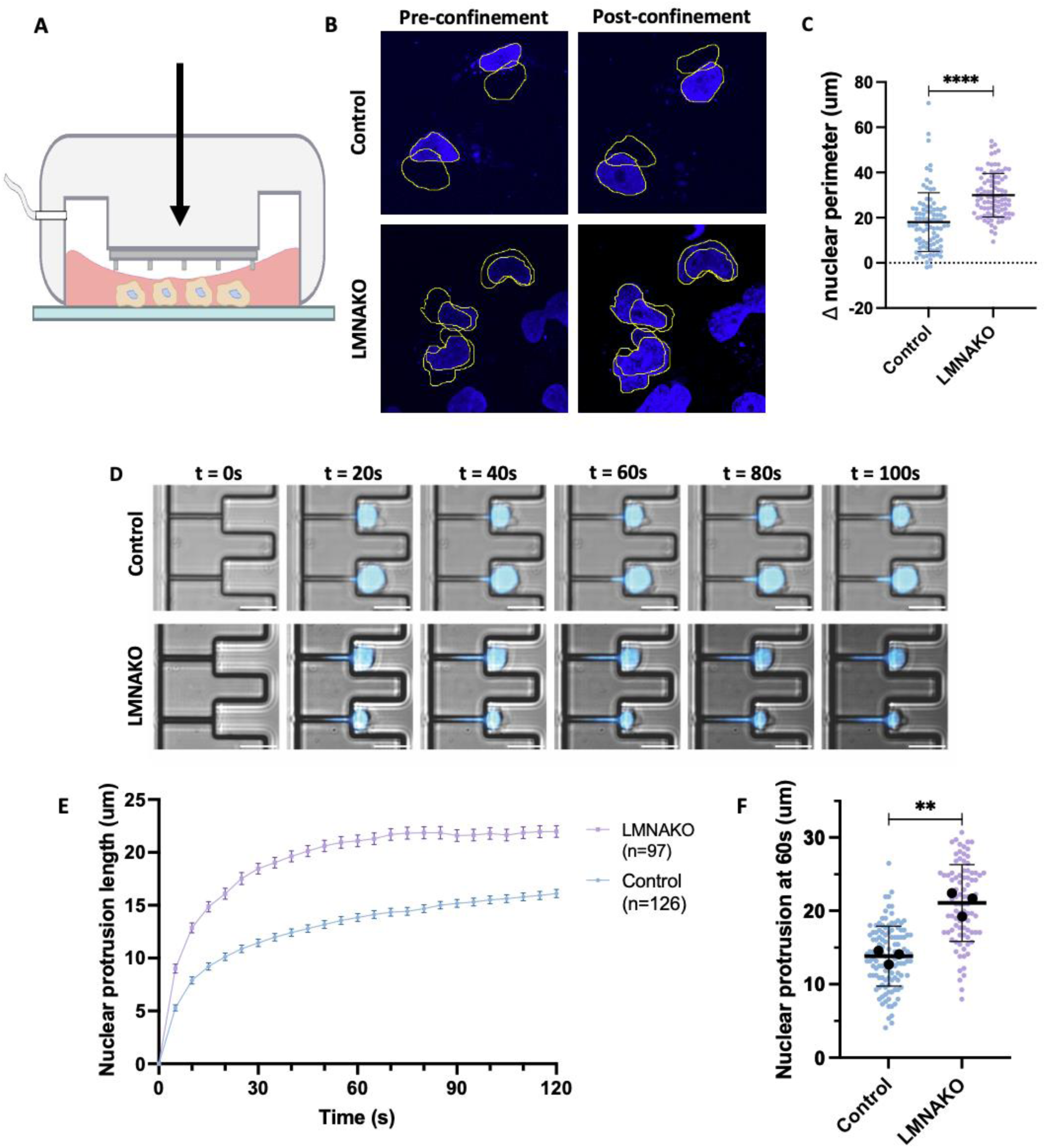
Lamin A loss increases nuclear deformability and migration through confined spaces. A: Schema of confinement assay B: Representative IF image with pairwise tracing of control and LMNAKO nuclei pre and post confinement C: Change in nuclear perimeter pre and post confinement D: Representative image of micropipette aspiration assay in DMS114 control and LMNA KO cells E: Nuclear protrusion of DMS114 control and LMNA KO over time, Errors bars are standard error of mean. F: Quantification of nuclear protrusion at t=60s Significance tests used parametric unpaired T-tests with Welch’s correction. **p<0.01, ****p<0.0001. Error bars are mean with standard deviation unless otherwise indicated.

To further test whether enhanced deformability in LMNAKO cells improves transit through confined spaces, we employed a microfluidic micropipette aspiration assay [80]. This assay confers large deformation on the nucleus in a manner that simulates in vivo pressurized passage through confined spaces such as narrow capillaries or interstitial spaces, as occurs in metastasis. LMNAKO nuclei were aspirated into small channels (3 × 5 μm² cross-section) more rapidly and further within 100 seconds than control nuclei (**Fig. 6D, E, F**, Movie S1, S2). Remarkably, 7 out of 97 LMNAKO cells fully perfused through a 30 μm channel within the time limit, whereas no control cells did (Movie S3), further underscoring the enhanced nuclear deformability of LMNAKO cells.

These results demonstrate that LMNA loss significantly increases nuclear deformability, facilitating the transit of SCLC cells through confined spaces. These findings align with existing literature emphasizing the importance of nuclear mechanics in cellular movement during processes such as extravasation and metastasis [76, 77, 81, 82]. Overall, our findings highlight the utility of our novel mouse model in elucidating the roles of specific genes in metastatic organotropism and suggest that low *LMNA* expression may enhance liver metastasis.

## Discussion

SCLC has remarkable metastatic proclivity, with most patients presenting with widely disseminated metastatic disease [6, 83]. SCLC exhibits a strong preference for metastasizing to specific organs, a phenomenon known as organotropism, with the liver and brain being the most common sites. However, our understanding of the mechanisms driving SCLC metastasis and the factors determining metastatic fate toward one organ over another has been limited by scarcity of metastatic samples and lack of robust pre-clinical models.

Here, we developed a series of SCLC xenograft models using tumors from the liver, lung, lymph node, and adrenal obtained through rapid autopsy. These models faithfully recapitulate key aspects of human SCLC, including organ colonization of the liver, brain, adrenal, bone marrow, and kidneys, as well as histopathological features and responses to clinically relevant therapies. A distinctive feature of our models is the rapid onset of metastatic lesions, occurring within weeks, mirroring the high metastatic propensity of SCLC in patients, which contrasts with metastasis models of other tumor types that typically require multiple rounds of *in vivo* selection [16–21]. Through *in vivo* lineage selection, we refined our models to achieve even shorter latency and improved penetrance, enabling us to identify cellular programs associated with organ-specific metastatic behaviors, particularly in the liver and brain. We expect these models to be valuable tools for investigating the pathways and factors that drive SCLC’s enhanced metastatic capabilities and organ-specific preferences.

Our findings reveal a consistent upregulation of stem-like pathways, especially the WNT signaling, alongside de-enrichment of cytoskeletal and nucleoskeletal organization in liver metastases. In contrast, brain metastases are characterized by the activation of pathways related to neuronal identity, cell-cell junctions, cytoskeletal remodeling, and extracellular matrix interactions. We validated the role of nuclear-cytoskeletal interactions, focusing on the nuclear envelope protein lamin A/C, encoded by the *LMNA* gene. Loss of *LMNA* increased nuclear deformability in vitro and significantly heightened the incidence of liver metastasis. We speculate that the dynamic regulation of *LMNA* expression, akin to mechanisms observed in extravasating neutrophils [84], may be a characteristic of cells that successfully metastasize, with the ability to downregulate LMNA levels being a key feature of metastatic potential. Moreover, LMNA loss may explain the well-documented phenomenon of nuclear molding in SCLC cells, where reduced cytoplasmic volume and distinct nuclear morphology are prominent diagnostic histological features.

Liver is the most frequent site of SCLC metastases, portending reduced responses to cancer therapies and dismal outcomes [85, 86]. Liver is also the most frequent site of metastasis across multiple cancer types [87]. Dissecting the molecular mechanisms of liver metastases has been challenging due to limited preclinical models that recapitulate this process. Notably, only three of over 100 established SCLC cell lines originate from liver metastases [8]. Our findings, which highlight the epigenetic and transcriptomic reprogramming in liver metastases that alter nucleoskeletal and cytoskeletal pathways, as well as the significant increase in metastatic burden following *LMNA* downregulation, underscore the importance of nuclear mechanics in liver metastasis [76, 77]. The patient-derived models developed in this study provide a valuable resource for further exploration of these mechanisms and for testing potential therapeutic interventions aimed at targeting the physical properties of cancer cells to prevent or reduce metastasis.

Brain metastases are a serious complication in SCLC, affecting 15-20% of patients at diagnosis and increasing to 40-60% as the disease progresses [14, 88]. Brain metastases are such a central clinical feature of SCLC that prophylactic cranial irradiation is standard care for SCLC patients [89]. However, our understanding of SCLC brain metastasis is limited by the scarcity of human brain metastasis samples and models. SCLC xenograft models rarely metastasize to the brain, instead developing leptomeningeal disease [90], and genetically engineered mouse models of SCLC exhibit a very low incidence of brain metastasis [91]. Recent studies of SCLC brain metastases used intracranial injection models, but these reveal narrow aspects of metastasis i.e. growth of an established metastatic tumor [42, 55]. A recent study utilizing our model demonstrated that vascular co-opting metastatic cells from brain metastasis models enter a state characterized by significant reduction in proliferation. This temporary state enables these cells to withstand various stressors, such as DNA damage, allowing them to survive and subsequently resume aggressive proliferation to colonize the brain. The upregulation of neuronal pathways observed in this context underscores the complex interplay between metastatic SCLC cells and the brain microenvironment. Overall, these findings underscore the relevance and validity of our model for exploring the mechanisms underlying SCLC brain metastasis.

While these models provide significant insights, they do not capture the whole metastatic cascade, which includes local invasion, intravasation, survival in transit, invasion, and colonization. Instead, they bypass the vascular invasion step. The observed decrease in 406B brain penetrance and number of liver metastases compared to past injection rounds, highlights the inherent variability of metastatic assays [92] despite rigorous procedural consistency. Additionally, these models lack an intact immune system, a significant limitation when examining the full spectrum of metastatic behavior. Another limitation is the focus on lamin A/C without distinguishing between its various isoforms. Future research will be necessary to explore the contributions of these isoforms to our findings based on the deletion of the *LMNA* gene. While we emphasized the mechanical roles of lamin A/C in this study, its contributions to epigenetic regulation may represent an additional mechanism that warrant further exploration. In conclusion, this study presents a novel preclinical model of SCLC organotropism and reveals the role of LMNA in liver metastasis, paving the way for the development of targeted therapies to prevent or treat metastatic disease.

## Methods

### Rapid Autopsy Tissue Processing

NIH IRB, Office of Human Subjects Research Protections at NCI, approved the studies; all patients provided written informed consent for tumor sample sequencing. The experiments conformed to the principles set out in the WMA Declaration of Helsinki and the Department of Health and Human Services Belmont Report. Normal and metastatic tissue samples were procured two hours after end of life. Fresh tissue samples were resected and divided into three pieces that were fixed in 10% formalin for paraffin embedding, flash frozen in liquid nitrogen, and/or stored fresh in MACS Tissue Storage Solution (Miltenyi Biotec 130-100-008) on ice for further manipulation. Fresh samples were rapidly transferred to lab and dissociated using Tumor Dissociation kit, human (Miltenyi Biotec 130-095-929) and gentleMACS Dissociator on 37°C hTDK-1 pre-set protocol. Dissociated cells were then used for scRNAseq when of high quality and to generate cell-lines by putting in RPMI-1640 supplemented with 10% FBS and 1% Pen-strep. Patient information was de-identified and all samples labeled dissection number-site of metastasis (eg: 6-liver)

### Mouse Tumor Model Generation

Animal studies were approved by the Animal Care and Use Committee (ACUC) of the NCI-Frederick. Frederick National Laboratory for Cancer Research (FNLCR) is accredited by AAALAC International and follows the Public Health Service Policy for the Care and Use of Laboratory Animals. Animal care was provided in accordance with the procedures outlined in the “Guide for Care and Use of Laboratory Animals (National Research Council; 1996; National Academy Press; Washington, D.C.).

*Four-six*<u>-week-old</u> male or female NSG mice (NOD.Cg-Prkdc scid Il2rg tm1Wjl/SzJ; # 005557, The Jackson Laboratory, Bar Harbor, ME) were injected intracardiac in the left ventricle with 500K tumor cells to generate each CDX model. Mice were evaluated weekly by MRI and BLI to assess tumor burden. In-vivo planar bioluminescence imaging was performed using IVIS Spectrum imaging system (Revvity, Waltham, MA) with the following specifications: excitation filter: block, emission filter: open, f/stop: 1, binning: medium (8X8) and exposure: auto (typically 1-120 seconds), mouse position: dorsal and ventral. Image acquisition and analysis were performed with vendor specific software LivingImage (version 4.7.4, Revvity, Waltham, MA). Bioluminescence signal to assess tumor burden in the liver was quantified by placing a fixed size rectangular region of interest (ROI) over the ventral image for each mouse. Similarly, tumor burden in the brain was assessed by analyzing the dorsal images with a fixed size circular ROI. Total flux (photons per second (s) per cm^2^ per steradian (sr) [photons/s/cm^2^/sr]) within each ROI was used as a comparing parameter. For drug treatment experiments, on day 5 following tumor cell injection, the mice were randomized based on bioluminescence flux signal and body weight using the StudyLog randomization software into treatment groups (n=14 for vehicles and n=5 per treatment group). The treatment regimen is listed in Table 1.

Mice were imaged weekly and weighed twice weekly. Mice were euthanized when the liver BLI flux signal exceeded 5×10^7^ or the brain BLI signal exceeded 1×10^7^ (photons per second (s) per cm2 per steradian (sr)), or body weights dropped below 20%. Drug doses and regimen were decided based on previously published studies [93–98]. For all non-drug experiments, 10 mice were injected per group and mice were euthanized as stated above.

### MRI method

Two weeks post-inoculation, MRI was initiated on the 3T clinical scanner (Intera-Achieva 3T, Philips, Netherlands), utilizing a custom-built volume receive array coil (Lambda Z Technologies, MD) for simultaneously imaging three mice. A scout sequence with slices in three orthogonal directions (sagittal, coronal, and axial) was employed to locate mouse anatomy and determine imaging planes.

A multi-slice T2-weighted turbo spin echo sequence (T2w-TSE) with a field of view (FOV) of 78 x 160 mm was applied in the coronal view from the brain to lower abdomen. The images were acquired with a repetition time (TR) of 5880 ms, an echo time (TE) of 45 ms, in-plane resolution of 0.18 × 0.18 mm², and a slice thickness of 0.5 mm.

A fat-suppressing technique, Spectral Presaturation with Inversion Recovery (SPIR), was employed to suppress fat components in the abdomen, thus improving tissue contrast and the visibility of metastatic lesions.

### CDX Tumor Processing

Mice were sacked and generation 1 and 2 BMD and LMD cell-lines were generated by resecting tumors, storing on ice in MACS Tissue Storage Solution (Miltenyi Biotec 130-100-008) until dissociation using Tumor Dissociation kit, human (Miltenyi Biotec 130-095-929) and gentleMACS Dissociator on 37°C hTDK-1 pre-set protocol. Select samples were treated with red blood cell lysis buffer before sending to the NCI Single Cell Analysis Facility for scRNA-seq (10X Genomics platform targeting 1000-6000 cells). Remaining cells were put in RPMI-1640 supplemented with 10% FBS and 1% Pen-strep to generate cell-lines.

### Cell Culture

All cell-lines culture in RPMI-1640 supplemented with 10% FBS and 1% Pen-strep at 37°C in a humidified atmosphere with 5% CO_2_. Luciferase tagged cell-lines were intermittently cultured in 1ug/ml Blasticidin (Gibco A1113903) and treated with luciferin and assessed for BLI using Avis to ensure it was maintained. 406B LMNA KO cell-line was cultured in media supplemented with 0.5ug/ml puromycin to maintain of the cas9 activity to prevent loss and potential for subsequent reversion mutations. Cell-lines were assessed routinely for mycoplasma contamination using the MycoAlert mycoplasma detection kit (Lonza).

### Generation of CRISPR/Cas9 constructs

Putative guide RNAs targeting protein coding exons were designed using sgRNA Scorer 2.0[99]. Five candidate guide RNAs (Table 2) were assessed for editing efficiency by generating in vitro transcribed (IVT) RNA and complexing with recombinant Cas9 protein and subsequently transfecting the complex into 293T cells. 48-72 hrs after transfection, DNA was extracted using Quick Extract and targeted amplicon sequencing was performed to quantify indel efficiencies, similar to what was done previously [100]. Oligonucleotides encoding candidates 712 (pCE0452) and 713 (pCE0453) were annealed and ligated into Lenti-CRISPR-V2 (Addgene# 52961) [101]. lentiCRISPR v2 was a gift from Feng Zhang (Addgene plasmid # 52961 ; http://n2t.net/addgene:52961 ; RRID:Addgene_52961). Constructs were then Sanger sequenced to verify successful guide RNA cloning. LMNA knockout constructs generated have been deposited in Addgene.

**Table 2.**
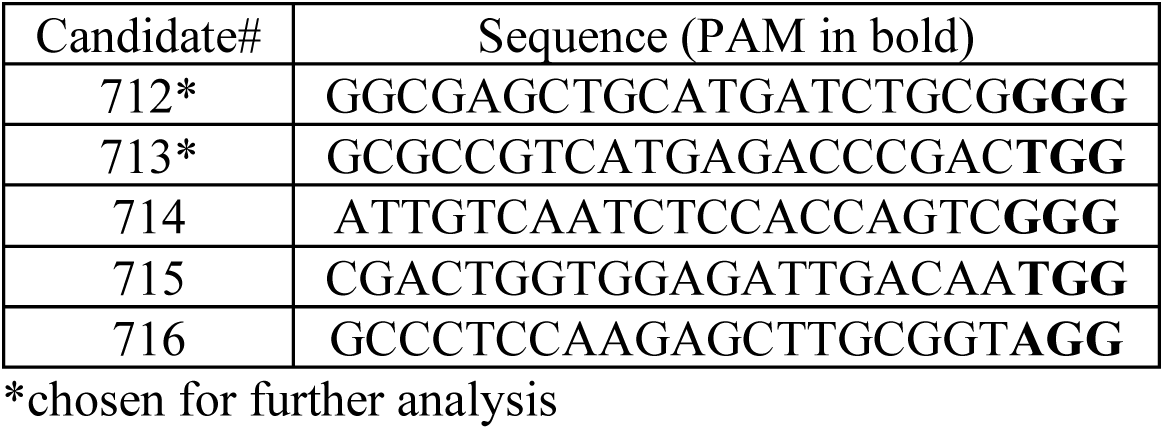
List of guide RNA sites designed and tested.

**Table 3.**
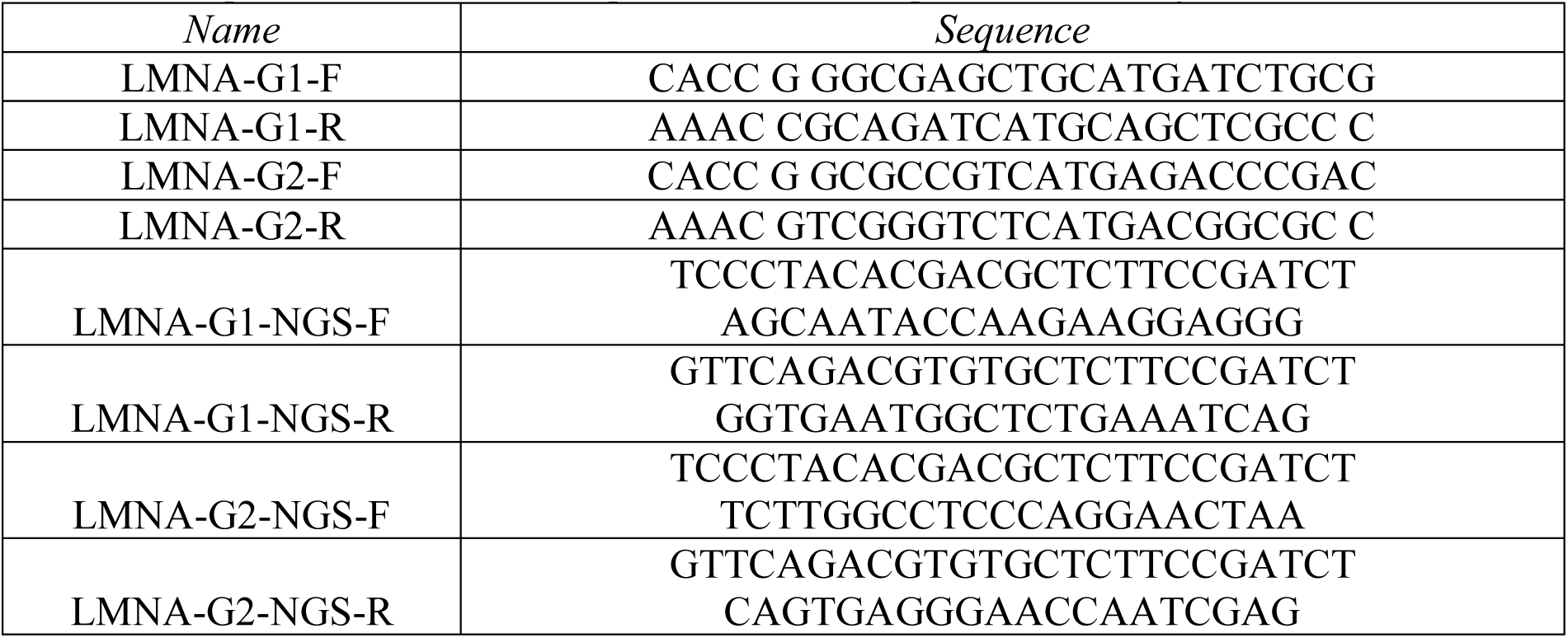
Oligonucleotides used for guide RNA cloning and clone analysis.

### Genetic modification of cell-lines

Lentiviruses were produced by transient transfection of HEK-293T cells with lentiviral plasmid of interest, Pax2 (Addgene 35002), and VSV.G (Addgene 14888). Lentiviral plasmids included pLenti CMV V5-LUC Blast (Addgene 21474) for luciferase addition and pCE0452 (Chari lab, NCI Fredrick) for LMNA KO. HEK-293T cells were seeded in DMEM +10% FBS on day -1, transfected using turbofectin 8.0 on day 0 (OriGene TF81001), and media containing virus was collected on days 1 and 2, purified, combined, and frozen for later usage. Infection was achieved by seeding 1-2 million of cell-line of interest in 1ml of viral soup. Cells were washed and put in RPMI 1647 +10% FBS +1% P/S and allowed to expand for two days before selection with 1ug/ml puromycin or 5ug/ml blasticidin for three days. For LMNA KO puromycin concentration was reduced to maintenance of 0.5ug/ml and cells allowed to further expand before single cell cloning for LMNA KO. For luciferase cells were reduced to a maintenance of 1ug/ml and eventually taken off blasticidin entirely. Maintenance of luciferase was monitored and occasionally cells put back on blasticidin to ensure no change. Negative controls were conducted for all transductions.

Single cell-cloning for LMNA KO was achieved by serial dilution into 96 well plates, visual assessment for wells that contain single cells, expansion over 1 month, followed by assessment for LMNA KO by western blot and DNA editing analysis using deep amplicon sequencing.

Briefly, 50 ng of genomic DNA extracted from knockout cells was used as template for PCR using the Q5 Ultra II 2X polymerase mastermix in a 20 uL reaction. PCR products were then sequenced on the Illumina MiSeq and raw fastq files were analyzed using the ngs_analysis_amplicon pipeline (https://github.com/rajchari2/ngs_amplicon_analysis). Sorted bam files were the visualized using the Integrative Genomics Viewer (IGV) [102]

### RNA-seq Collection, Dataset Preprocessing and Bioinformatics Analysis

RA22-6 and derived cell-lines cultured were collected (∼1 million per tube), washed with PBS, and RNA Isolated using the RNeasy plus mini kit (Qiagen 74134) to collect total RNA. Three technical replicates were used from each cell line. RNA integrity was verified using a standard sensitivity assay (Agilent 4150 TapeStation system). Samples were then shipped to Novogene for mRNA library preparation (polyA enrichment) and NovaSeq PE 150 (6 G raw data per sample) sequencing. Upon receipt of data, the paired-end RNA-seq fastq files were aligned with the hg38 human reference genome. STAR aligner (version 2.7.11b) and SAMtools software (version 1.17) used to generate sorted and indexed BAM files. ‘–quantMode GeneCounts’ module of STAR aligner utilized to compute the read counts per gene for RNA-seq datasets.

Subsequently, the raw read counts were normalized to RPKM (Reads Per Kilobase per Million mapped reads) values. After that, differential expression analysis was performed within different metastatic samples, using DEseq2 package in R (version 4.2.3). The significantly up-regulated (log2FC > 1.5) and down-regulated (log2FC < -1.5) genes with p-value (< 0.05) were considered for the pathway analysis. Ingenuity Pathway Analysis (IPA, http://www.ingenuity.com) software tool used for the identification of important biological pathways [38]. In this analysis, we used the cutoffs -1 >z-score >1 and -log10 p-value >1.3 and then ordered by z-score to select differentially enriched pathways.

### Assay of Transposase Accessible Chromatin sequencing

Cell-lines were frozen at ∼2 million cells per tube and shipped to novogene for library preparation and ATAC-seq by NovaSeq PE150 (9 G raw data per sample). Upon receipt of data, we check the sequencing quality of reads using FASTQC software (version 0.12.1) and the duplicate reads were detected and filtered out using Picard software (version 3.2.0). BWA (version 0.7.17) aligner software used to generate aligned BAM files with hg38 reference genome. MACS2 “callpeak” module (version 2.2.7.1) used to generate peak calling .BED files. BAMscale “scale” (version 0.0.6) module used to generate normalized bigwig files[103]. We have used the “cov” module of the BAMscale software to calculate genome-wide normalized FPKM coverages of each BAM files. Bedtools “subtract” (version 2.31.1) and deepTools (version 3.5.5) “plotHeatmap” module used to detect the unique peaks and to generate average signal intensity heatmaps with in the parent and different generations metastatic cell lines. To discover enriched motifs in these unique peak regions, we have employed findMotifsGenome.pl module of HOMER software with default parameters. It provides list of motifs, consensus sequence, p-value, -log_10_(p-value), q-value, % target sequence with motif and % background sequence with motif. For further analysis, we have selected top-ranked motifs based on the -log_10_(p-value) for parent and gen1/gen2 metastatic samples. In addition, “limma” package use to identify differentially enriched peaks and Genomic Regions Enrichment of Annotations Tool (GREAT) version 4.0.4 used to identify pathways within different generations and parent. Pathways were curated based on RegionFoldEnrich > 1.5, ObsGenes >9, and then selected and grouped based on relevance. Redundant pathways based on gene similarity were excluded from presentation in figures. IGV (Integrative Genomics Viewer) tool (version 2.16.0) used for the visualization of ATAC signals at various genomic coordinates. All the volcano plots, Principle component analysis (PCA) plots, dot plots and correlation heatmaps were generated using “matplotlib.pyplot” and “seaborn.clustermap” and “sklearn.decomposition” python (version 3.9.17) libraries.

### Multiome sequencing

Optimal lysis time was first determined by nuclei isolation using different lysis times and assessment of nuclei integrity with widefield microscope. A lysis time of 3 minutes and 30 seconds was selected. On the day, 10X Genomics nuclei isolation single cell multiome ATAC + gene expression sequencing protocol was followed: https://cdn.10xgenomics.com/image/upload/v1660261285/support-documents/CG000365_DemonstratedProtocol_NucleiIsolation_ATAC_GEX_Sequencing_RevC.pdf. Briefly, 1 million cells were used per sample, lysis and washes conducted, and then cell aliquots were counted using propium iodide (Thermo Scientific Invitrogen Countess 3 Automated Cell Counter) and aliquots assessed by immunofluorescence (Zeiss LSM 780) with DAPI staining for nuclei integrity assessed. Nuclei samples were next transposed using 10x Genomics Chromium Next GEM Single Cell Multiome ATAC + Gene Expression User Guide. The transposed nuclei were loaded in the lanes with one capture lane per sample targeting recovery around 6,000 nuclei per lane. Partitioning was completed successfully with uniform emulsion consistency. Reverse transcription and barcoding were followed immediately after the partitioning. All subsequent steps of library preparation and quality control were performed as described in the 10x user guide for Multiome ATAC + GEX.

Next, filtered feature-barcode matrices for RNA and ATAC-seq data were loaded using Read10X_h5. A Seurat object was created for each dataset, with RNA counts set as the default assay and ATAC peaks with specified fragments. The annotations were derived from the UCSC hg38 genome. Three datasets (reference: RA22-6, test: Gen1BMD, Gen1LMD) were merged into a single Seurat object. For ATAC data, peaks across datasets were reduced to those with widths between 20 and 10,000 bp, and merged counts were obtained. Cells across all datasets were filtered based on RNA (nFeature_RNA > 2500 & < 7500, percent.mt < 30) and ATAC metrics (nFeature_ATAC > 1000 & < 30000, TSS.enrichment > 1, nucleosome_signal < 2).

Normalization and Dimensionality Reduction: RNA counts were normalized using SCTransform, and RNA-based dimensionality reduction was performed via RunPCA and RunUMAP (dims = 1:40). ATAC features were processed with RunTFIDF, RunSVD, and RunUMAP (dims = 2:40). Multimodal neighbors were identified with FindMultiModalNeighbors, integrating RNA (PCA) and ATAC (LSI) embeddings.

Clustering, Differential Expression and Motif Enrichment: Clusters were identified using FindClusters (resolution = 0.2) based on weighted nearest neighbors. Differentially expressed genes (DEGs) were identified using FindAllMarkers (test.use = “wilcox”, min.pct = 0.1, abs(avg_log2FC) > 1, p_val_adj < 0.01) for both RNA and ATAC modalities. Marker peaks in ATAC were linked to genes using LinkPeaks. Motif enrichment was performed using FindMotifs on ATAC peaks specific to clusters. Motif annotations were derived from the JASPAR2020 database, and p-values were adjusted using the Benjamini-Hochberg method.

Pathway Enrichment and Gene Imputation Analysis: Pathway enrichment analysis was performed using the AUCell framework on SCT-normalized gene expression values. Pathway activity scores for each cell were extracted from the AUC matrix and used for downstream analyses. MAGIC (Markov Affinity-based Graph Imputation of Cells) imputation was applied to smooth single-cell gene expression data. Specific genes of interest were filtered from the MAGIC-imputed matrix and exported for further validation and visualization. All analyses were conducted in R using the Seurat and Signac packages. Custom code in python was utilized for creating specific charts.

### IHC and H&E Staining

IHC staining was performed on LeicaBiosystems’ BondRX autostainer with the following conditions: Epitope Retrieval 1 (Citrate) 20’ for Synaptophysin, c-myc, Ki67, Chromogranin A, Lamin A sections, Epitope Retrieval 2 (EDTA) 10’ for Thyroid Transcription Factor TTF-1 sections, Epitope Retrieval 2 (EDTA) 20’ for CD56 NCAM1 sections, Epitope Retrieval 2 (EDTA) 25’ for INSM1 sections, Synaptophysin (abcam #32127, 1:1600 30’), c-myc (abcam #ab32072, 1:40 30’), Ki67 (Cell Signaling Technology #9027, 1:200 30’), Chromogranin A (Novus Biologicals #NB120-15160, 1:3000 30’), Lamin A (Cell Signaling Technology #86846, 1:500 30’), TTF-1 (abcam #ab72876, 1:100 30’), CD56 (Cell Signaling Technology #99746, 1:100 30’), INSM1 (Santa Cruz #sc-271408, 1:1000 30’), and the Bond Polymer Refine Detection Kit (LeicaBiosystems #DS9800). Isotype control reagents were used in place of primary antibodies for the negative controls. Slides were removed from the Bond autostainer, dehydrated through ethanols, cleared with xylenes, and coverslipped.

H&E slides were stained on the Tissue-Tek® Prisma™ autostainer.

### Western Blotting

Cells were collected, protein lysates isolated using lysis buffer + proteinase inhibitor cocktail (Invitrogen # 1861281), and concentrations assessed and normalized using a BCA assay. NuPAGE™ LDS Sample Buffer (Invitrogen, # 1879570) + sample reducing agent (Invitrogen, lot 1658579) was then added to 1X working concentration and denatured at 95°C for 5 minutes. Denatured proteins were separated on a precast NuPAGE bis-tris Gel and transferred to a PVDF transfer stack (thermofisher) using iBlot 2 system. Dry transfer (Thermofisher iBlot2) was conducted at 20V for 6min. The membrane was then blocked for 1hr in 5% skim milk in TBST and incubated in primary mouse monoclonal antibody Lamin A/C 4C11 (Cell Signaling Technology #4777) in 5% BSA overnight at 4°C. Goat anti-mouse HRP-conjugated secondary antibody (Invitrogen 31430) was used to visualize using chemiluminescence buffers (Invitrogen 34580) and the Biorad ChemiDoc Imaging System.

### DNA FISH

Metaphase spread was performed as described https://ccr.cancer.gov/sites/default/files/metaphase_preparation_from_adherent_cells.pdf.Cells were imaged using a Leica Thunder Imager.

### Confinement assays

Cellular confinement during high-resolution time lapse live-cell imaging was performed on a Nikon Eclipse Ti2 microscope equipped with a Yokogawa CSU-W1 scanhead (Nikon) and a motorized stage with a Nano-Z100 piezo insert (Mad City) and a Plan Apo 60x oil 1.49 NA objective. Illumination was provided by the Nikon LUNV 6-line laser unit and captured with a Hamamatsu Orca-Flash 4.0 v3 camera. The system was controlled with Nis-Elements (Nikon). Confinement was induced with the 1-well Dynamic Cell Confiner System (4Dcell) utilizing a cobalt autonomous vacuum pump attached to a PDMS cup fitted with a glass coverslip containing PDMS micropillars 3 μm in height. Prior to confinement experiments, DMS114 cells were plated for 24 hours on 35mm glass bottom dishes (FluoroDish, WPI) coated with 10 μg/ml fibronectin in RPMI media. The PDMS cup and coverslip were briefly sonicated in 70% ethanol, followed by PBS (Invitrogen), then placed in cell culture media for 1 hour at room temp to equilibrate the PDMS. To confine the cells, the 35 mm glass bottom dish was pre-incubated for 30 min with 1 ml of media containing Hoechst 33342 (1:1000 ThermoFisher) to label the nuclei, then washed out prior to imaging. Utilizing a low pressure vacuum seal the PDMS was attached to the dish and confinement was initiated over 2 min using a ramp of 10 mbar per second.

Images were acquired from multiple XY positions every 2 min pre and post confinement to ensure full unbiased coverage of the confined area over a 1hr time period.

### Micropipette Aspiration Assay

Micropipette aspiration was performed using a previously published protocol [80]. In brief, a cell suspension was perfused into the micropipette device under defined pressure. The pressure at the top inlet port was set to 1 psi, and the bottom port was set to 0.2 psi. Cells were suspended in a buffer containing 2% BSA, 0.2% FBS, and 10 mM EDTA in PBS. Hoechst 33342 was added 1:1000 immediately before the cell suspension was transferred to devices. After establishing a flow of cells into the device, pockets were cleared to allow fresh cells to begin aspiration, and images were acquired every 5 s for 40 frames using a Zeiss Observer Z1 epifluorescence microscope with Hamamatsu Orca Flash 4.0 camera. For the micropipette experiments, we used a 20X LD Plan-Neoflaur Air Objective; NA = 0.4. The image acquisition for micropipette aspiration experiments was automated with Zen Blue (Zeiss) software.

## Acknowledgements

This project has been funded in whole or in part with Federal funds from the National Cancer Institute, National Institutes of Health, under Contract No. HHSN261201500003I. The content of this publication does not necessarily reflect the views or policies of the Department of Health and Human Services, nor does mention of trade names, commercial products, or organizations imply endorsement by the U.S. Government.” This contract number represents work performed within the scope of work of the non-severable IDIQ contract.

## Data Availability

RNA-seq, ATAC-seq, sc-RNA-ATAC-seq data will be made available

