## Supplementary Data for "Metastatic organotropism in small cell lung cancer"

**Grant information:** This study was supported by the Center for Cancer Research, the Intramural Program of the NCI (ZIA BC 011793). A.T: grants to NCI from EMD Serono Research & Development, AstraZeneca, Gilead Sciences, and ProLynx. JL: NIH R35 GM153257, NSF URoL-2022048, Volkwagen Foundation A130142

Table S1- Cell-line information

| Cell line | Procurement method | Site of metastasis | Generation | Cell line injected to yield | Generation + site name used throughout paper |
|---|---|---|---|---|---|
| 4-lymph node | Human rapid autopsy | Cervical lymph node | Parental | | |
| 5-liver | Human rapid autopsy | Right lobe posterior inferior liver | Parental | | |
| 6-liver | Human rapid autopsy | Right lobe inferior most | Parental | | Parent |
| 12-lung | Human rapid autopsy | | Parental | | |
| 18-adrenal | Human rapid autopsy | Left adrenal | Parental | | |
| 400L | Mouse necropsy | Liver | 1 | 6-liver | Gen1LMD |
| 404L | Mouse necropsy | Liver | 1 | 6-liver | Gen1LMD |
| 406B | Mouse necropsy | Brain | 1 | 6-liver | Gen1BMD |
| 408B | Mouse necropsy | Brain | 1 | 6-liver | Gen1BMD |
| 431L | Mouse necropsy | Liver | 2 | 406B | Gen2LMD |
| 431B | Mouse necropsy | Brain | 2 | 406B | Gen2BMD |
| 438L | Mouse necropsy | Liver | 2 | 406B | Gen2LMD |
| 438B | Mouse necropsy | Brain | 2 | 406B | Gen2BMD |

Table S2- Immunohistochemical staining of SCLC markers

| Tumor Sample | Target | Marker Type | Typical [1] | IHC Type | Total Cells | % Positive | H-Score |
|---|---|---|---|---|---|---|---|
| RA-22-6 | Chromogranin A | Traditional NE Marker | (74% cases) | Cytoplasmic | 9111 | 89.5 | 210.3 |
| RA-22-6 | Synaptophysin | Traditional NE Marker | some not all cells | Cytoplasmic | 15830 | 99.1 | 293.4 |
| RA-22-6 | INSM1 | NE Marker | 92% | Nuclear | 74025 | 87.9 | 144.4 |
| RA-22-6 | CD56 | Traditional NE Marker | 75-100%(>80% of cases) | Cytoplasmic | 27528 | 99.5 | 298.0 |
| RA-22-6 | Ki-67 | Cancer Proliferative activity | 60% | Nuclear | 25155 | 58.7 | 93.2 |
| 406-Liver | Chromogranin A | Traditional NE Marker | (74% cases) | Cytoplasmic | 5472 | 81.5 | 149.4 |

| | | | | | | | |
|---|---|---|---|---|---|---|---|
| 406-Liver | Synaptophysin | Traditional NE Marker | some not all cells | Cytoplasmic | 3033 | 99.2 | 296.5 |
| 406-Liver | INSM1 | NE Marker | 92% | Nuclear | 3671 | 74.3 | 87.6 |
| 406-Liver | CD56 | Traditional NE Marker | 75-100%(>80% of cases) | Cytoplasmic | 2201 | 99.9 | 299.4 |
| 406-Liver | Ki-67 | Cancer Proliferative activity | 60% | Nuclear | 2495 | 42.4 | 52.5 |
| 406-Brain | Chromogranin A | Traditional NE Marker | (74% cases) | Cytoplasmic | 1396 | 48.7 | 75.9 |
| 406-Brain | Synaptophysin | Traditional NE Marker | some not all cells | Cytoplasmic | 1786 | 99.7 | 298.7 |
| 406-Brain | INSM1 | NE Marker | 92% | Nuclear | 2079 | 89.2 | 124.1 |
| 406-Brain | CD56 | Traditional NE Marker | 75-100%(>80% of cases) | Cytoplasmic | 1801 | 99.8 | 297.6 |
| 406-Brain | Ki-67 | Cancer Proliferative activity | 60% | Nuclear | 17530 | 58.1 | 86.0 |

Table S3- Epigenetically linked differentially expressed genes in each cluster

| **Upregulated genes** | | | | | | **Downregulated genes** | | | | |
|---|---|---|---|---|---|---|---|---|---|---|
| **Cluster 1** | **Cluster 2** | **Cluster 3** | **Cluster 5** | **Cluster 0** | | **Cluster 0** | | | | |
| PPARGC1A | CDH1 | C1orf21 | SOX3 | L3MBTL4 | DMKN | SLC35F4 | GRIN2B | NRXN3 | VAV3 | ACVR1C |
| | ADGRD2 | CCDC152 | ARHGEF9 | MET | YBX3 | LINC01194 | NPAS3 | DHRS2 | SLC43A3 | LINC00461 |
| | CST4 | CACNG4 | PKD1L3 | NTS | ATP2B3 | ADARB2 | FRMD4B | LRRTM3 | MARCH4 | MAPK10 |
| | CD163L1 | SAMD13 | MYO5B | DSP | TRPM3 | LINC01470 | ZNF91 | TRHDE | RAB27B | |
| | | CTDSPL | COL4A4 | TNS3 | OPRD1 | BCL11B | PREX2 | NMUR2 | CRIM1 | |
| | | RHOJ | COL4A3 | PAPPA | PRPH2 | SEMA3A | PCDH17 | LIN7A | EYA2 | |
| | | LINC01151 | TGFBR3 | SLC18A1 | NR2F1 | KCNIP4 | AIG1 | STMN2 | COL4A6 | |
| | | NRXN2 | PTH2R | CNTN1 | ARHGAP22 | DSCAM | SKAP1 | SNTB1 | STC1 | |
| | | GADL1 | CR1L | RASSF9 | PKHD1 | ZNF536 | CTNNA3 | RAMP1 | NKD1 | |
| | | SPAG17 | | PRSS12 | ZNF486 | CADM2 | CNTNAP4 | PPP2R2B | UGT2A1 | |

| | | | | | | | | | |
|---|---|---|---|---|---|---|---|---|---|
| | | ZBTB7C | | LGR6 | MIR100HG | GABBR2 | GRIK1 | ARID5B | LINC00626 |
| | | KCTD16 | | ZNF90 | PDZRN4 | AMPH | DISC1FP1 | AFF3 | RNF152 |
| | | | | PPP4R4 | CHL1 | CST1 | CLMP | CACNA1E | SLC8A3 |
| | | | | | ERVMER61-1 | FGF13 | ZEB2 | PDE9A | CELF5 |

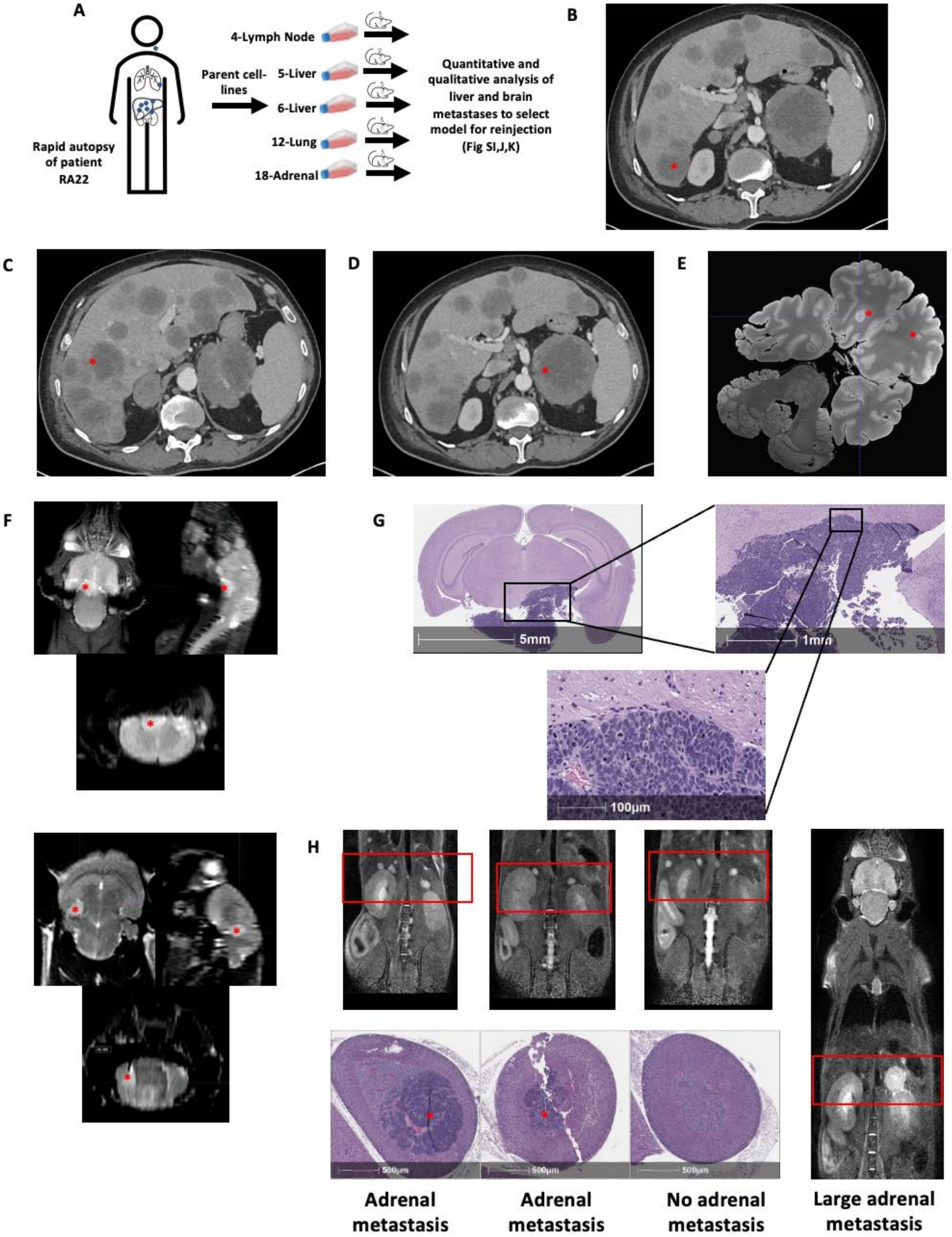

A
4-Lymph Node
Parent cell-lines
5-Liver
6-Liver
12-Lung
18-Adrenal
Rapid autopsy of patient RA22
Quantitative and qualitative analysis of liver and brain metastases to select model for reinjection (Fig SI,J,K)
B
C
D
E
F
G
5mm
1mm
100µm
H
500µm
Adrenal metastasis
Adrenal metastasis
No adrenal metastasis
Large adrenal metastasis

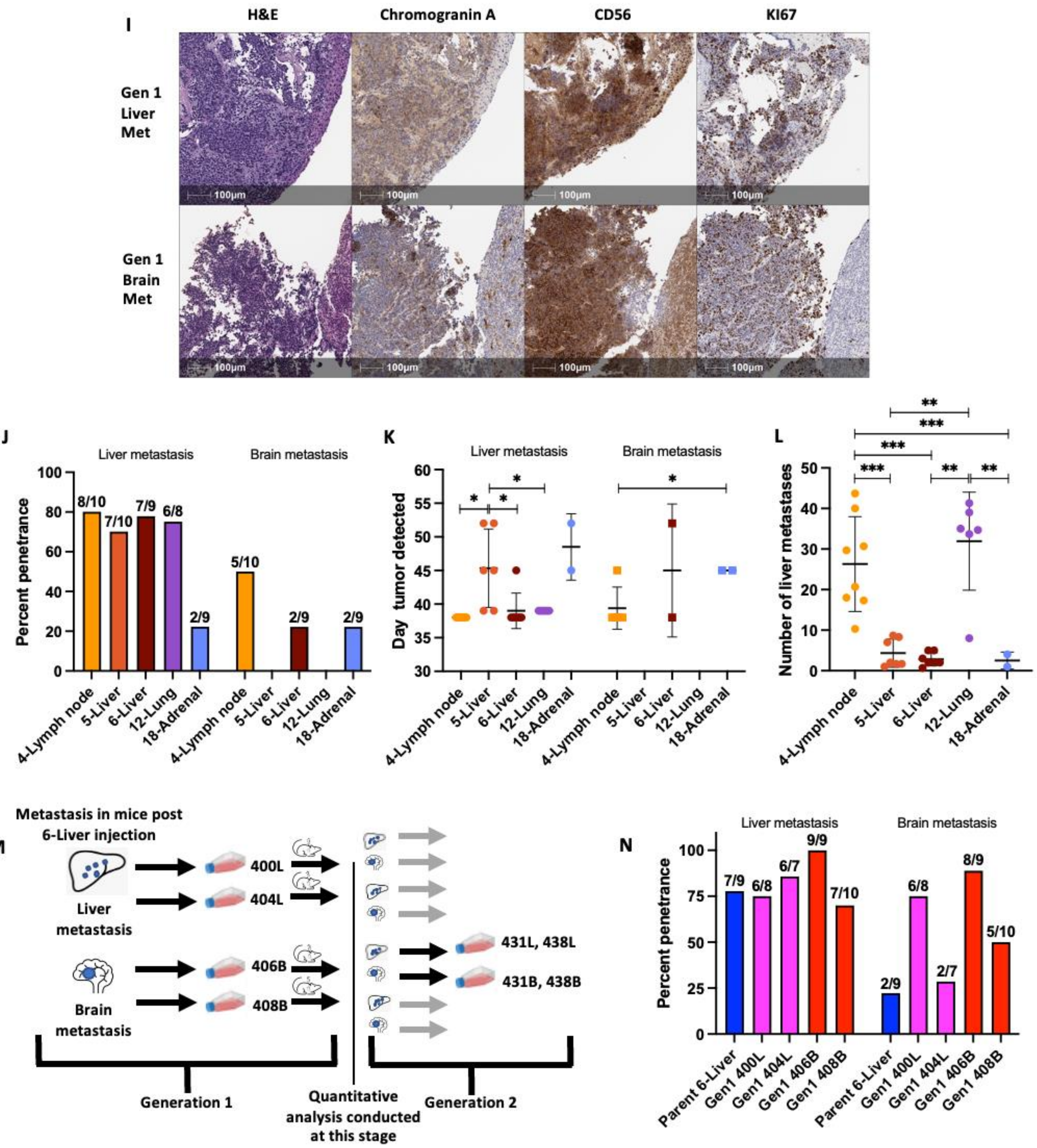

I
H&E
Chromogranin A
CD56
KI67
Gen 1 Liver Met
Gen 1 Brain Met
100µm
J
Liver metastasis
Brain metastasis
Percent penetrance
8/10
7/10
7/9
6/8
2/9
5/10
2/9
2/9
4-Lymph node
5-Liver
6-Liver
12-Lung
18-Adrenal
K
Liver metastasis
Brain metastasis
Day tumor detected
L
Number of liver metastases
M
Metastasis in mice post 6-Liver injection
Liver metastasis
Brain metastasis
400L
404L
406B
408B
431L, 438L
431B, 438B
Generation 1
Quantitative analysis conducted at this stage
Generation 2
N
Liver metastasis
Brain metastasis
Percent penetrance
7/9
6/8
6/7
9/9
7/10
2/9
6/8
2/7
8/9
5/10
Parent 6-Liver
Gen1 400L
Gen1 404L
Gen1 406B
Gen1 408B

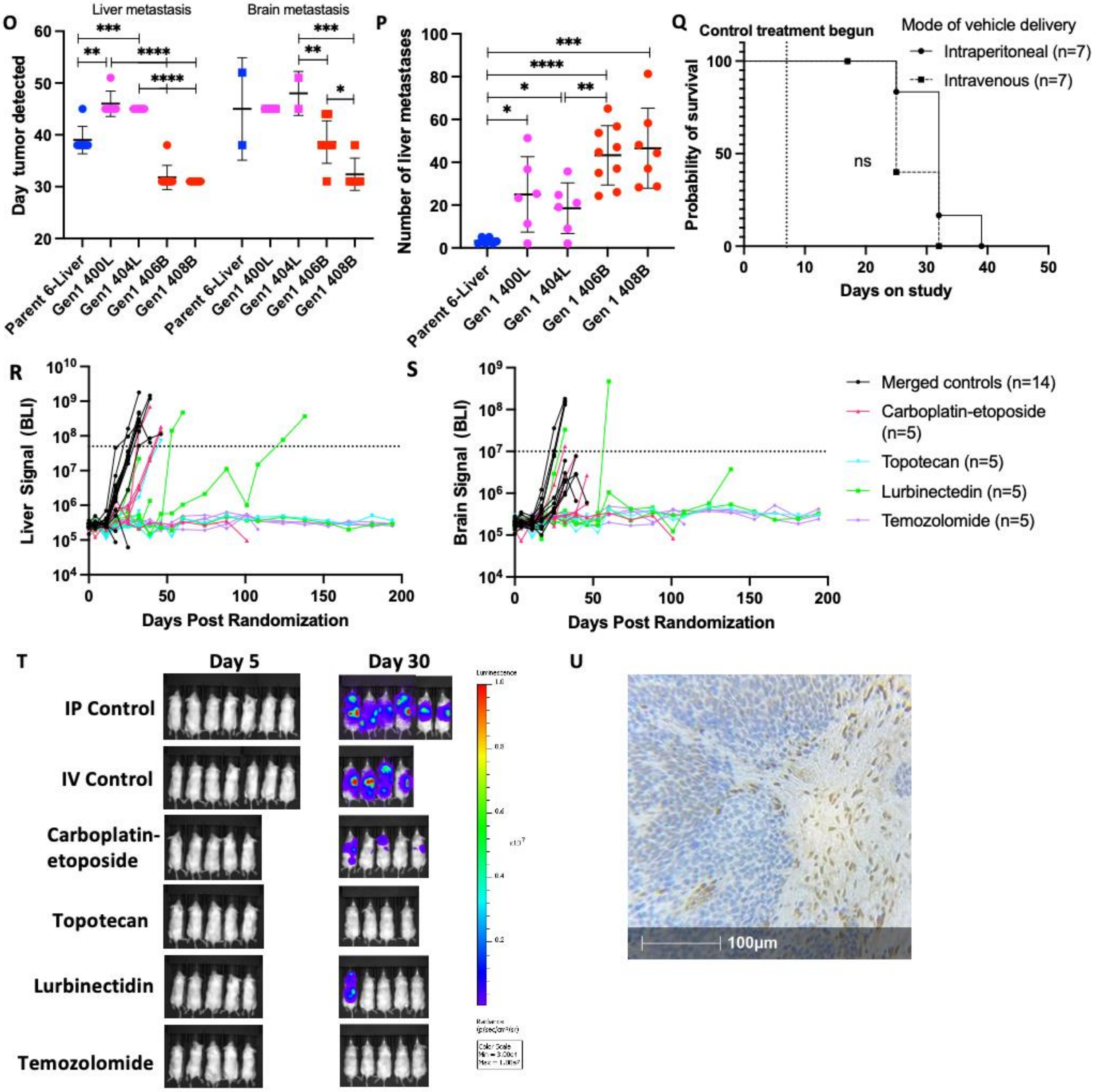

Figure S1: Establishment and characterization of a patient-derived model of SCLC metastases and organ tropism

A: Schema of mouse experiments with five parental cell lines.

B: CT scan for patient RA22 showing right posterior inferior lobe liver metastasis used to make 5-liver cell line, asterisks indicate metastasis

C: CT scan for patient RA22 showing right most inferior lobe liver metastasis used to make 6-liver cell line, asterisks indicate metastasis

D: CT scan for patient RA22 showing left adrenal metastasis used to make 18-adrenal cell line, asterisks indicate metastasis

E: Brain, CT post-mortem, asterisks indicate metastasis

F: Coronal, sagittal, and transverse cuts of typical (top) and rare (bottom) route of brain colonization, MRI, after RA22-4 (lymph node derived) cell line injection

G: Zoom of brain metastasis cells in Fig. 1F

H: Complexity identifying adrenal metastases on MRI

I: H&E and IHC for lung specific, tumor specific, and additional neuroendocrine factors in generation 1 metastases.

J: Percent penetrance (% of mice that formed tumors) of brain and liver metastases, assessed by MRI. RA22-4, RA22-5, RA22-6, RA22-12, RA22-18. Mice excluded if found dead or sacked due to bodyweight loss prior to appearance of first met in cohort. Mice numbers indicated above each bar.
K: Time to develop liver and brain metastasis in days. Mice with no metastases excluded.
L: Number of liver metastases. Mice with no metastases excluded
M: Schema of mouse experiments with five parental cell lines. RNA-seq data collected for generation 1 and 2 cell lines labeled.
N: Percent penetrance (% of mice that formed tumors) of brain and liver metastases, assessed by MRI. RA22-6, 400L, 404L, 406B, 408B. Mice excluded if found dead or sacked due to bodyweight loss prior to appearance of first met in cohort. Mice numbers indicated above each bar.
O: Time to develop liver and brain metastasis in days. Mice with no metastases excluded.
P: Number of liver metastases. Mice with no metastases excluded
Q: Survival (Kaplan-Meyer) curve of mice injected with 406B cells + vehicle. Vehicle delivered by routes indicated.
R: BLI signal from liver. The dotted horizontal line is the threshold for high tumor burden.
S: BLI signal from brain. The dotted horizontal line is the threshold for high tumor burden.
T: Representative image of BLI for each treatment condition
U: MGMT IHC from patient RA22 right most inferior lobe liver metastasis (used to make 6-liver cell line)

All significance indicated is following parametric unpaired T-tests with Welch's correction. *p<0.05, **p<0.01, ***p<0.001, ****p<0.0001. All error bars are mean with standard deviation unless mentioned otherwise. BMD: Brain metastasis derived; LMD cell lines: 400L, 404L; BMD cell lines: 406B, 408B. All patient Computational Topography (CT) transverse slices 2.5 weeks pre-mortem unless otherwise indicated.

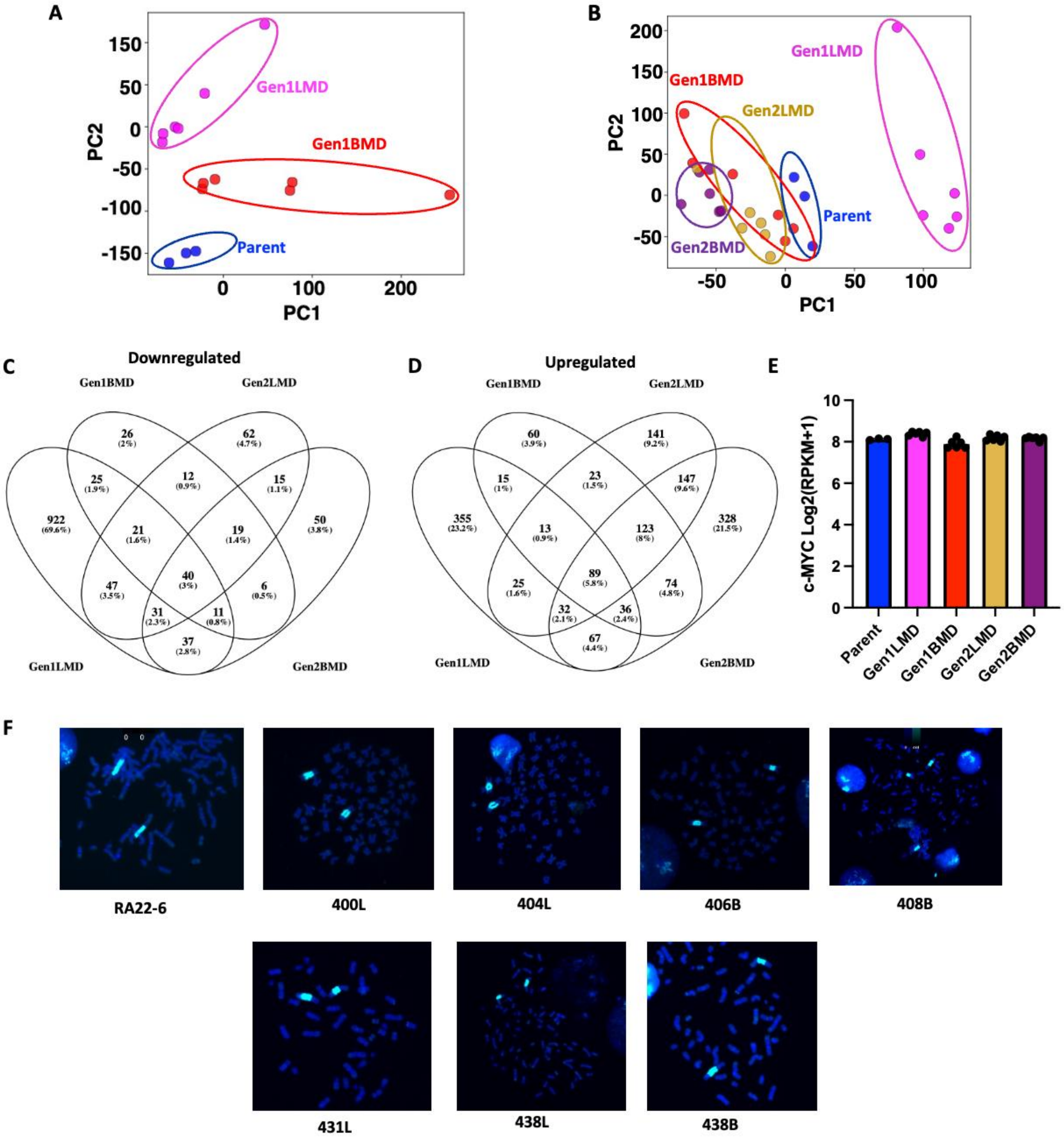

**Figure S2: Global gene expression analysis of RA22-6 and derived cell lines**

A: PCA of Parent and generation 1 all genes RNA-seq

B: PCA of Parent, generation 1, and generation 2 all genes RNA-seq

C: Venn diagram depicting differentially (Pvalue <0.05 & abs(logFC) >1.5) downregulated genes in Gen1LMD, Gen2LMD, Gen1BMD, Gen2BMD compared to Parent.

D: Venn diagram depicting differentially (Pvalue <0.05 & abs(logFC) >1.5) upregulated genes in Gen1LMD, Gen2LMD, Gen1BMD, Gen2BMD compared to Parent.

E: Gene expression of MYC

F: MYC DNA-FISH for MYC in Parent RA22-6, generation 1, and generation 2 cell lines

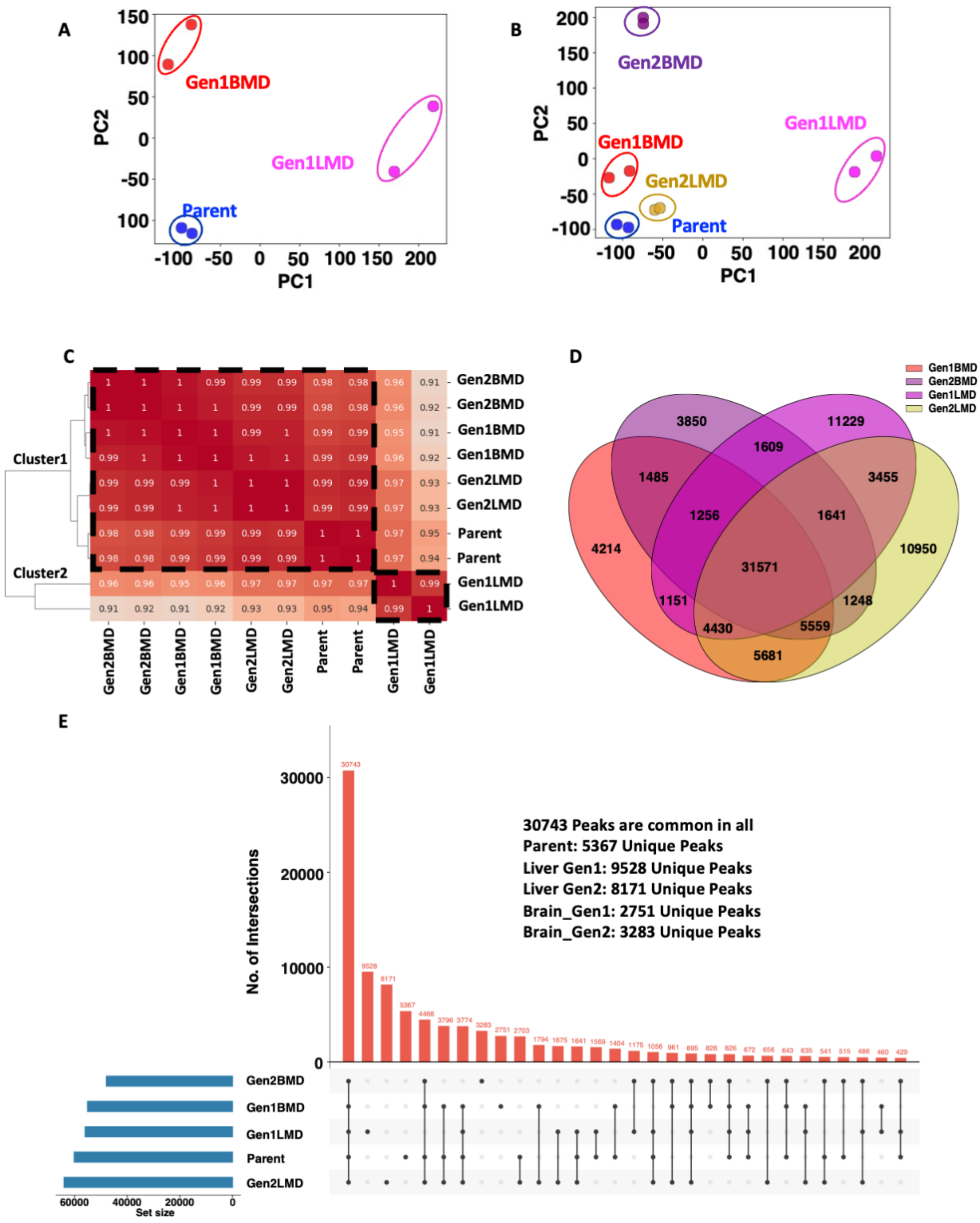

**Figure S3: Chromatin accessibility associated with SCLC organ-specific metastatic behaviors**

A: PCA of Parent and generation 1 all peaks ATAC-seq

B: PCA of Parent, generation 1, and generation 2 all peaks ATAC-seq
C: Pairwise correlation heatmap of open peaks in Parent, generation 1, and generation 2.
D: Venn Diagram depicting unique peaks between Gen1LMD, Gen1BMD, Gen2LMD and Gen2BMD.
E: Bar graph depicting overlap of peaks between Parent, generation 1, and generation 2 all peaks

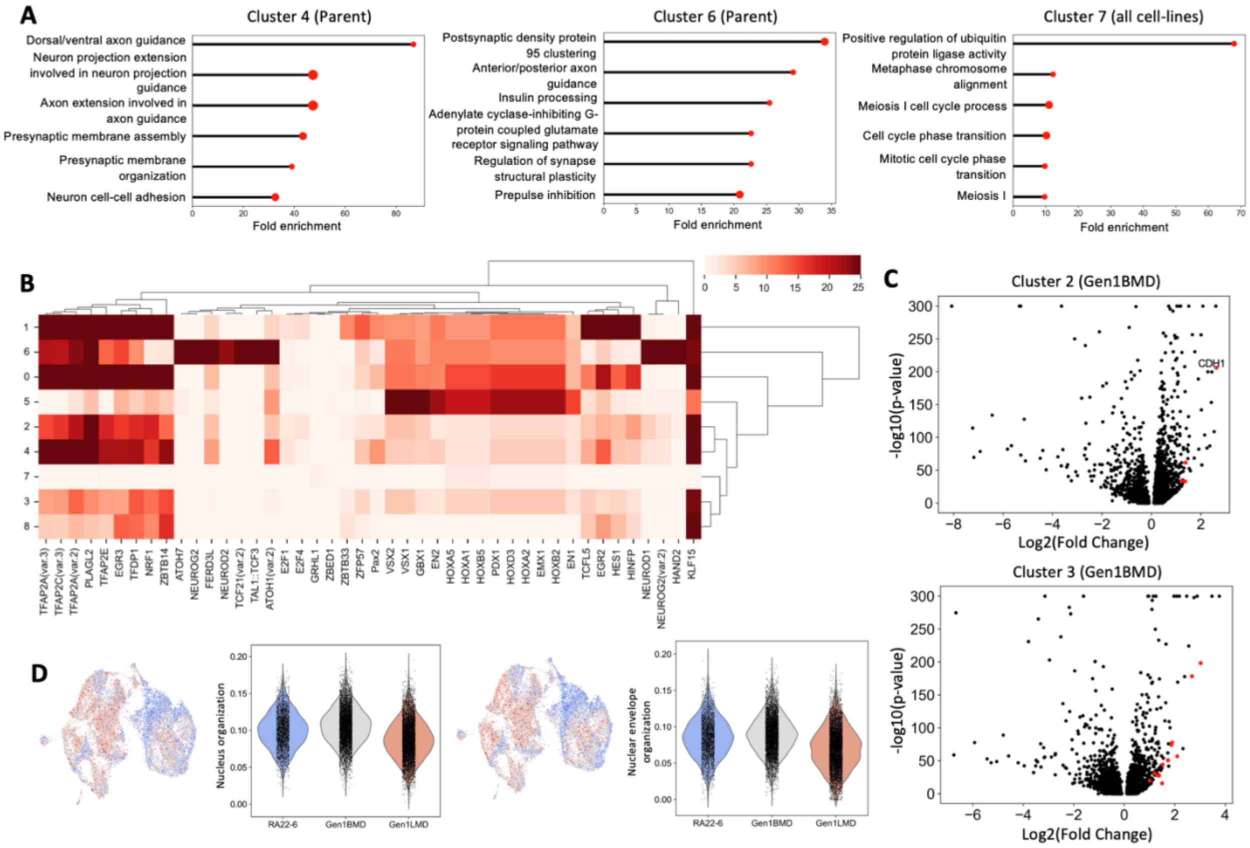

**E**

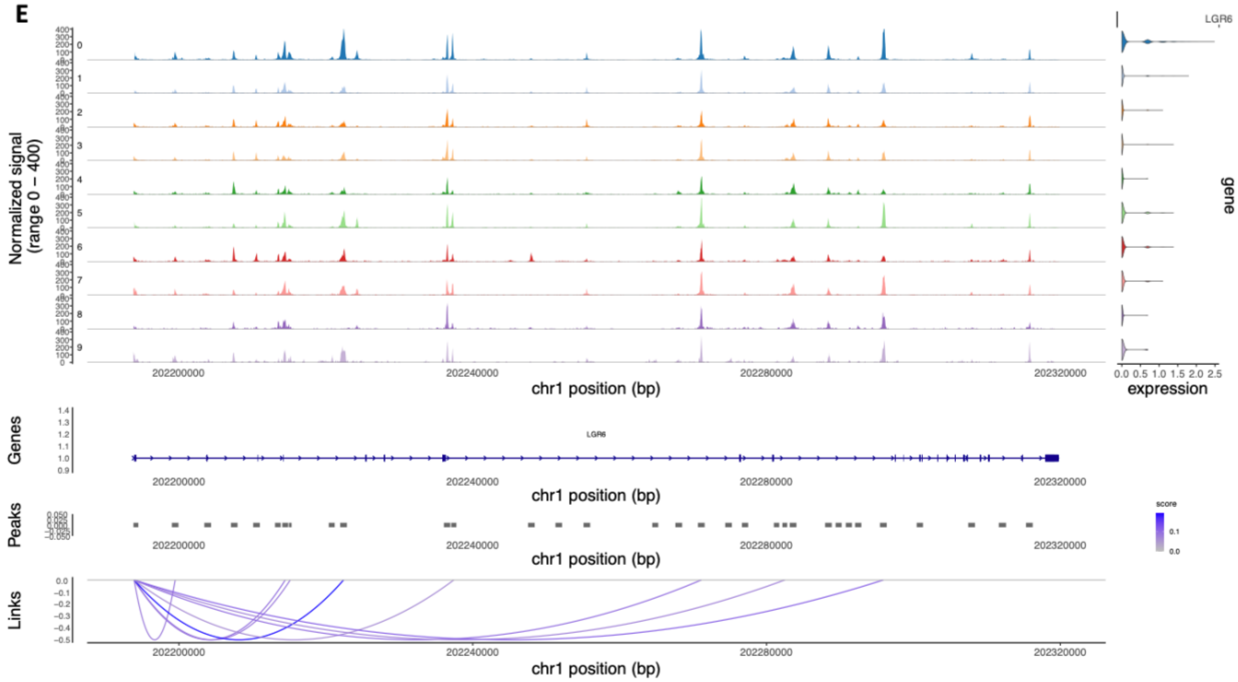
Normalized signal
(range 0 – 400)
LGR6
gene
expression
0.0 0.5 1.0 1.5 2.0 2.5
202200000
202240000
202280000
202320000
chr1 position (bp)
Genes
LGR6
Peaks
score
Links

**F**

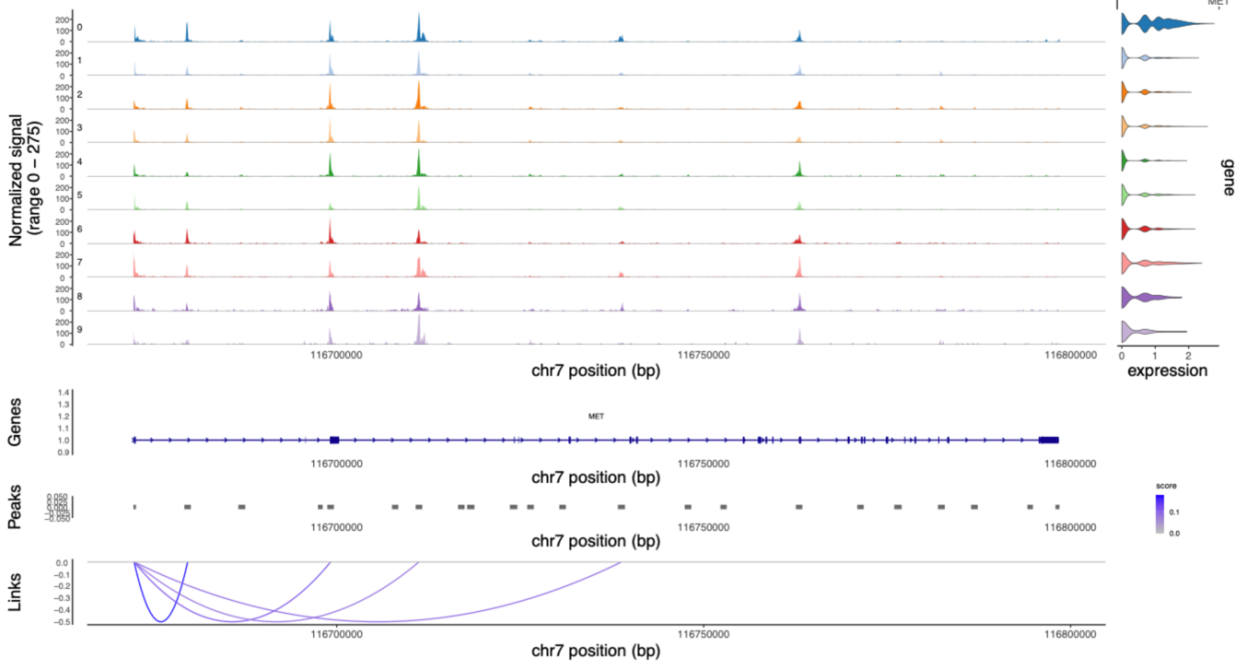
Normalized signal
(range 0 – 275)
MET
gene
expression
0 1 2
116700000
116750000
116800000
chr7 position (bp)
Genes
MET
Peaks
score
Links

G

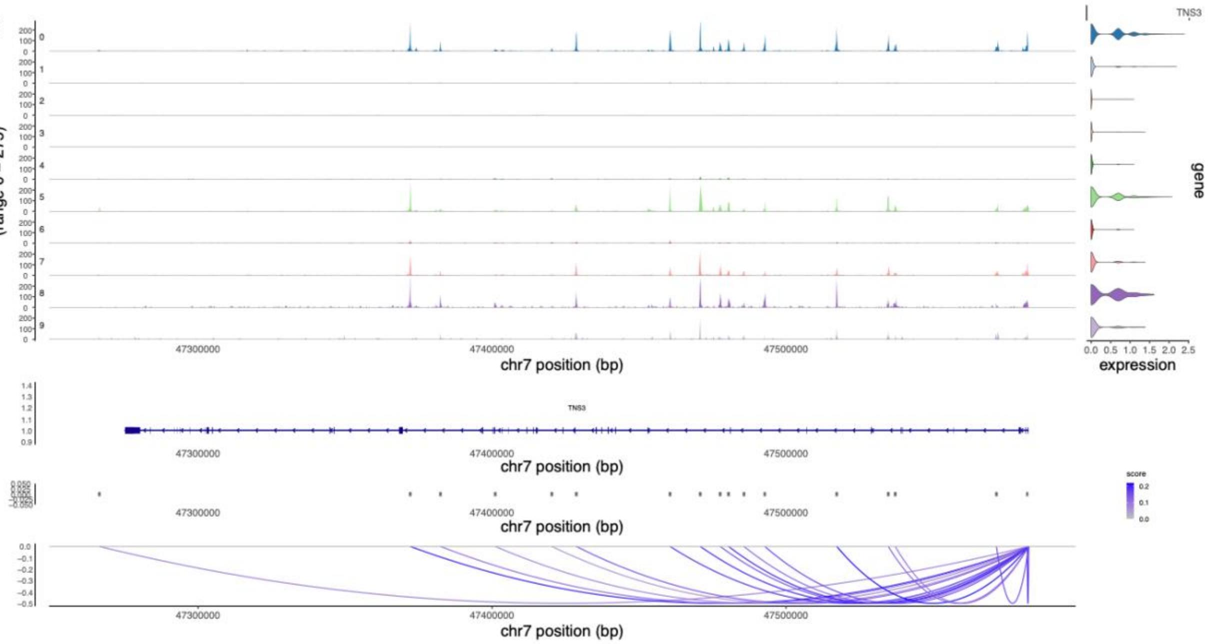

TNS3
Normalized signal
(range 0 – 275)
0
1
2
3
4
5
6
7
8
9
47300000
47400000
47500000
chr7 position (bp)
gene
0.0 0.5 1.0 1.5 2.0 2.5
expression
Genes
TNS3
Peaks
score
0.2
0.1
0.0
Links

H

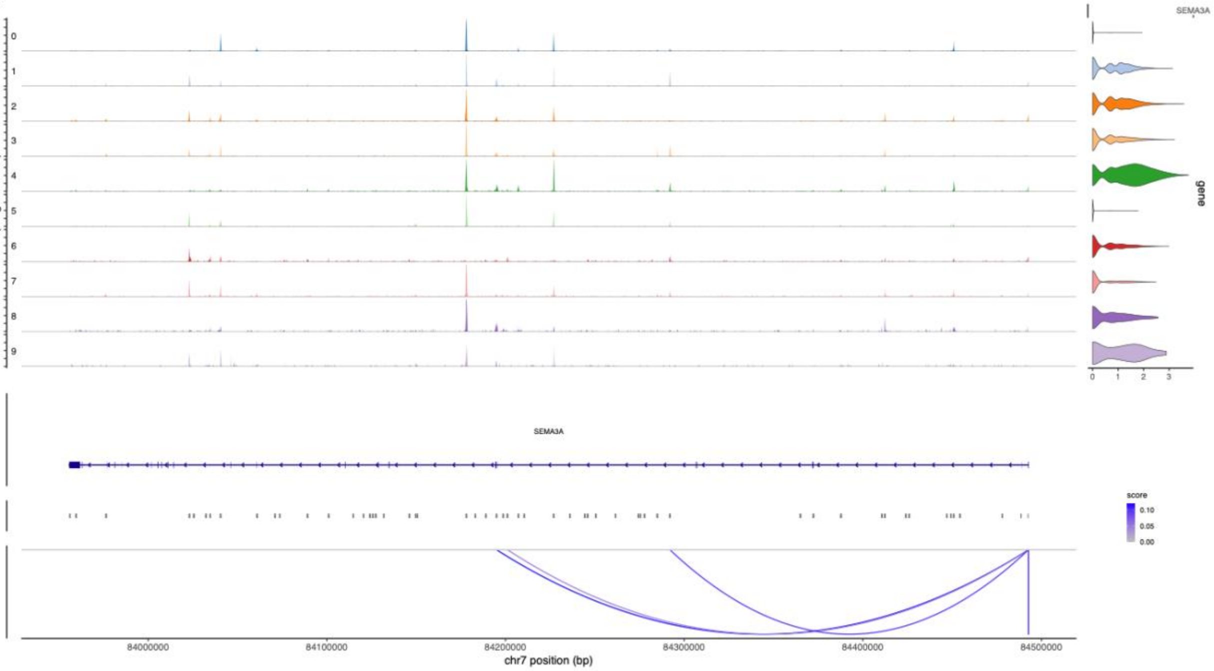

SEMA3A
Normalized signal
(range 0 – 200)
0
1
2
3
4
5
6
7
8
9
gene
0 1 2 3
Genes
SEMA3A
Peaks
score
0.10
0.05
0.00
Links
84000000
84100000
84200000
84300000
84400000
84500000
chr7 position (bp)

I

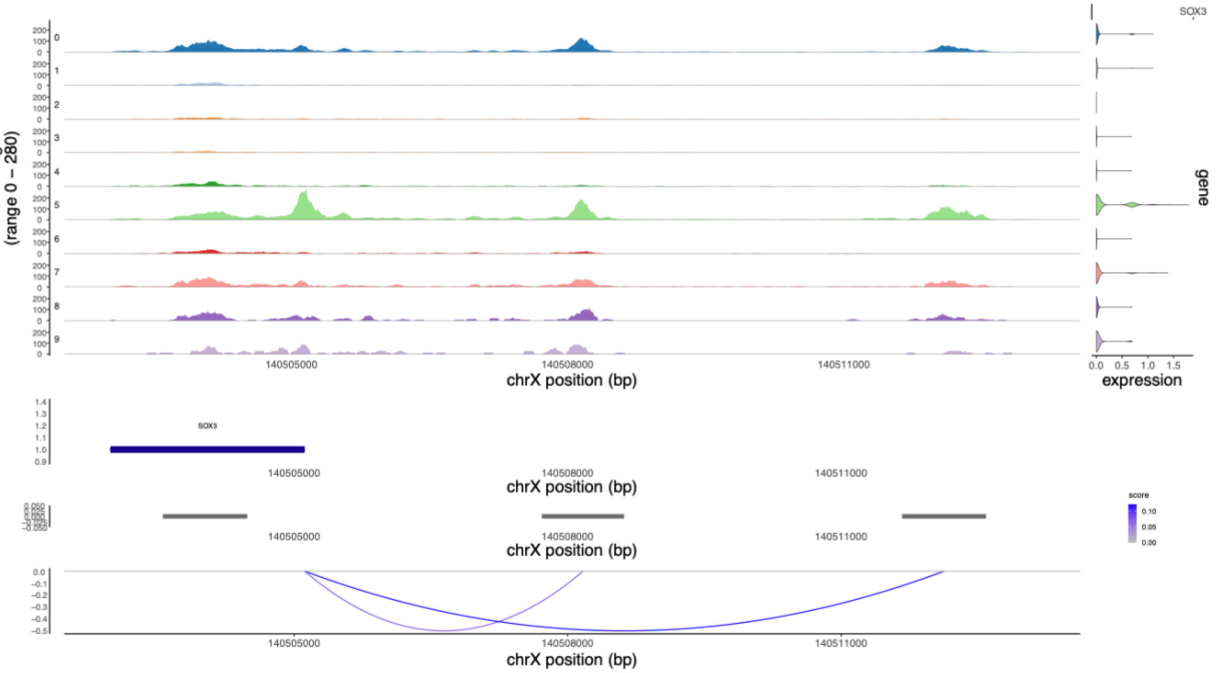

SOX3
Normalized signal (range 0 – 280)
0
1
2
3
4
5
6
7
8
9
200
100
0
140505000
140508000
140511000
chrX position (bp)
gene
0.0 0.5 1.0 1.5
expression
Genes
SOX3
1.4
1.3
1.2
1.1
1.0
0.9
Peaks
score
0.10
0.05
0.00
Links
0.0
-0.1
-0.2
-0.3
-0.4
-0.5

J

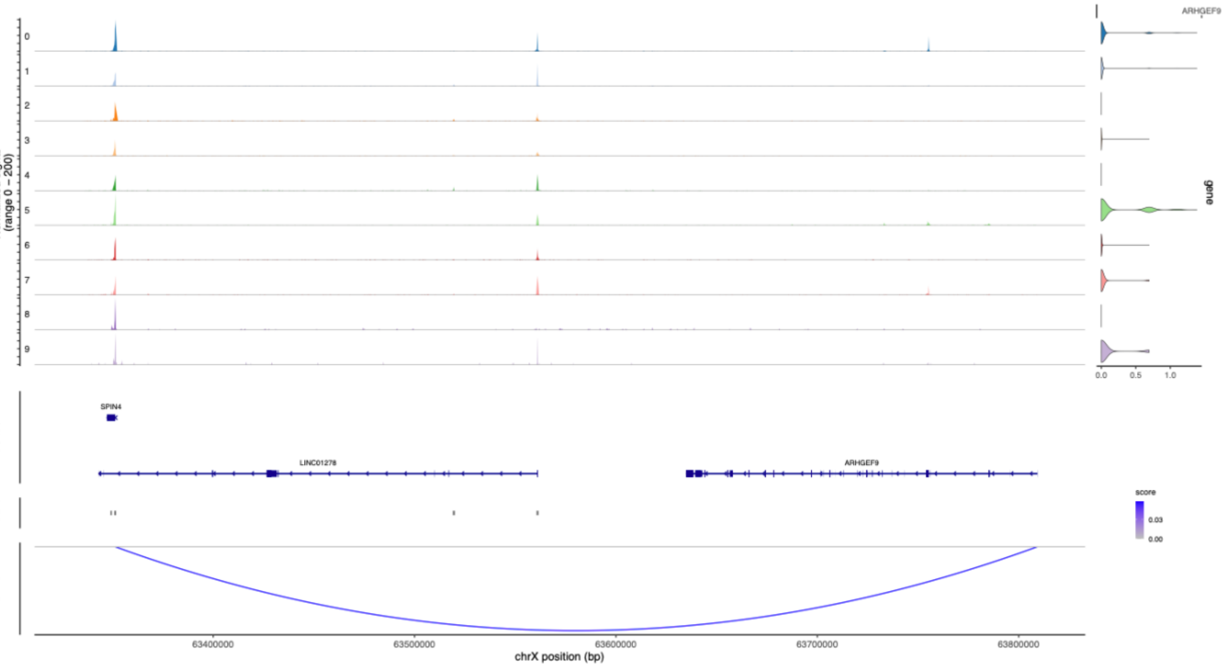

ARHGEF9
Normalized signal (range 0 – 200)
0
1
2
3
4
5
6
7
8
9
gene
0.0 0.5 1.0
Genes
SPIN4
LINC01278
ARHGEF9
Peaks
score
0.03
0.00
Links
63400000
63500000
63600000
63700000
63800000
chrX position (bp)

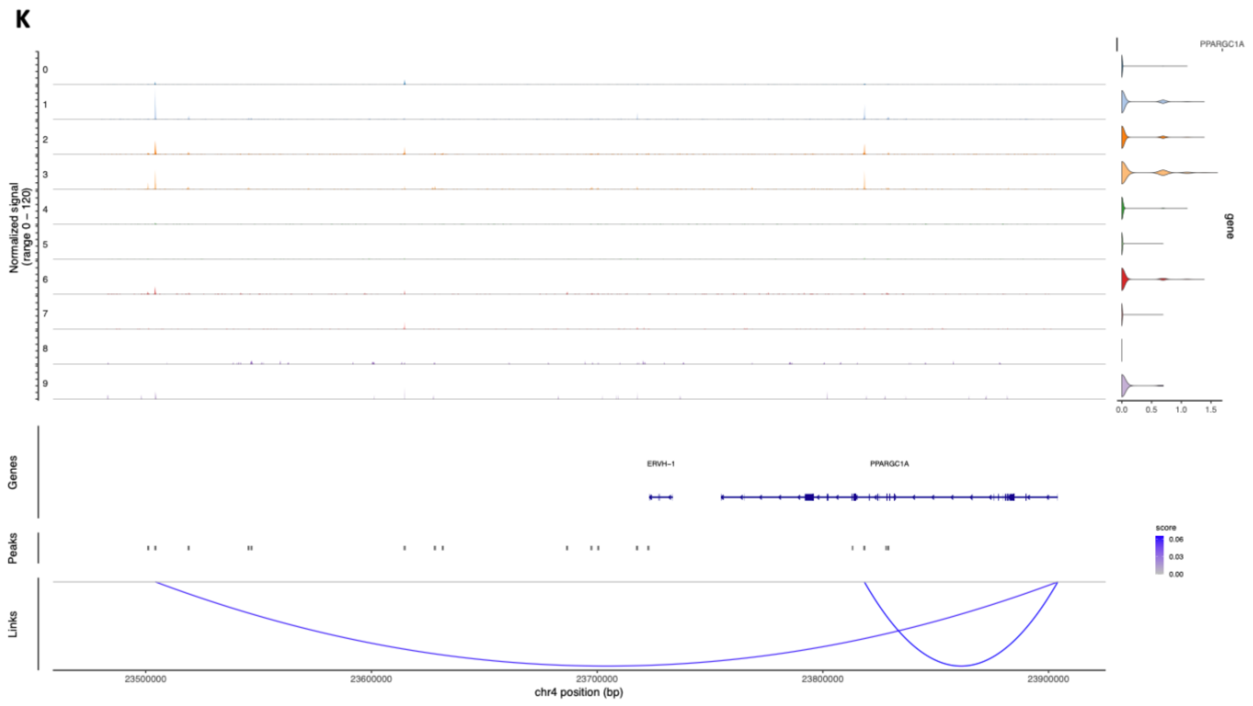

**Figure S4: Multiome sequencing supporting data**

A: Gene ontology of Parent (Clusters 4.6) and all samples (cluster 7).

B: Top 10 most significant known motifs for all clusters (44 motifs represented on graph as there was overlap). Complete list of motifs in supplemental file.

C: GSEA of nucleosome envelop organization and nucleosome organization from bulk ATAC-seq

D: Volcano plots of differentially expressed genes in individual clusters compared to all others. Red dots indicate genes that also show significant epigenetic linkage.

E: LGR6 coverage plot.

F: MET coverage plot.

G: TNS3 coverage plot.

H: SEMA3A coverage plot.

I: SOX3 coverage plot.

J: ARHGEF9 coverage plot.

K: PPARGC1A coverage plot.

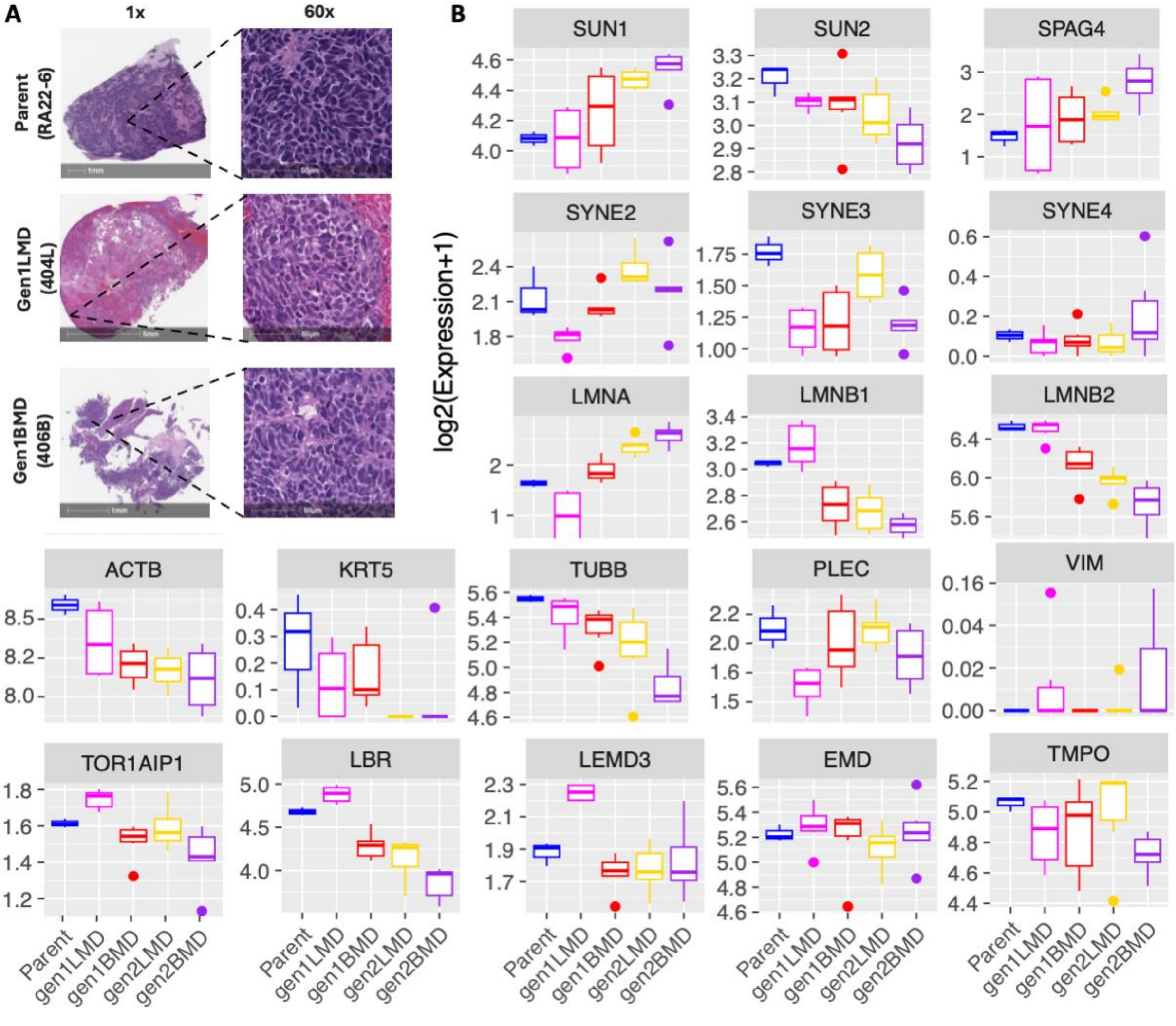
A
1x
60x
Parent (RA22-6)
Gen1LMD (404L)
Gen1BMD (406B)
B
log2(Expression+1)
SUN1
SUN2
SPAG4
SYNE2
SYNE3
SYNE4
LMNA
LMNB1
LMNB2
ACTB
KRT5
TUBB
PLEC
VIM
TOR1AIP1
LBR
LEMD3
EMD
TMPO
Parent
gen1LMD
gen1BMD
gen2LMD
gen2BMD

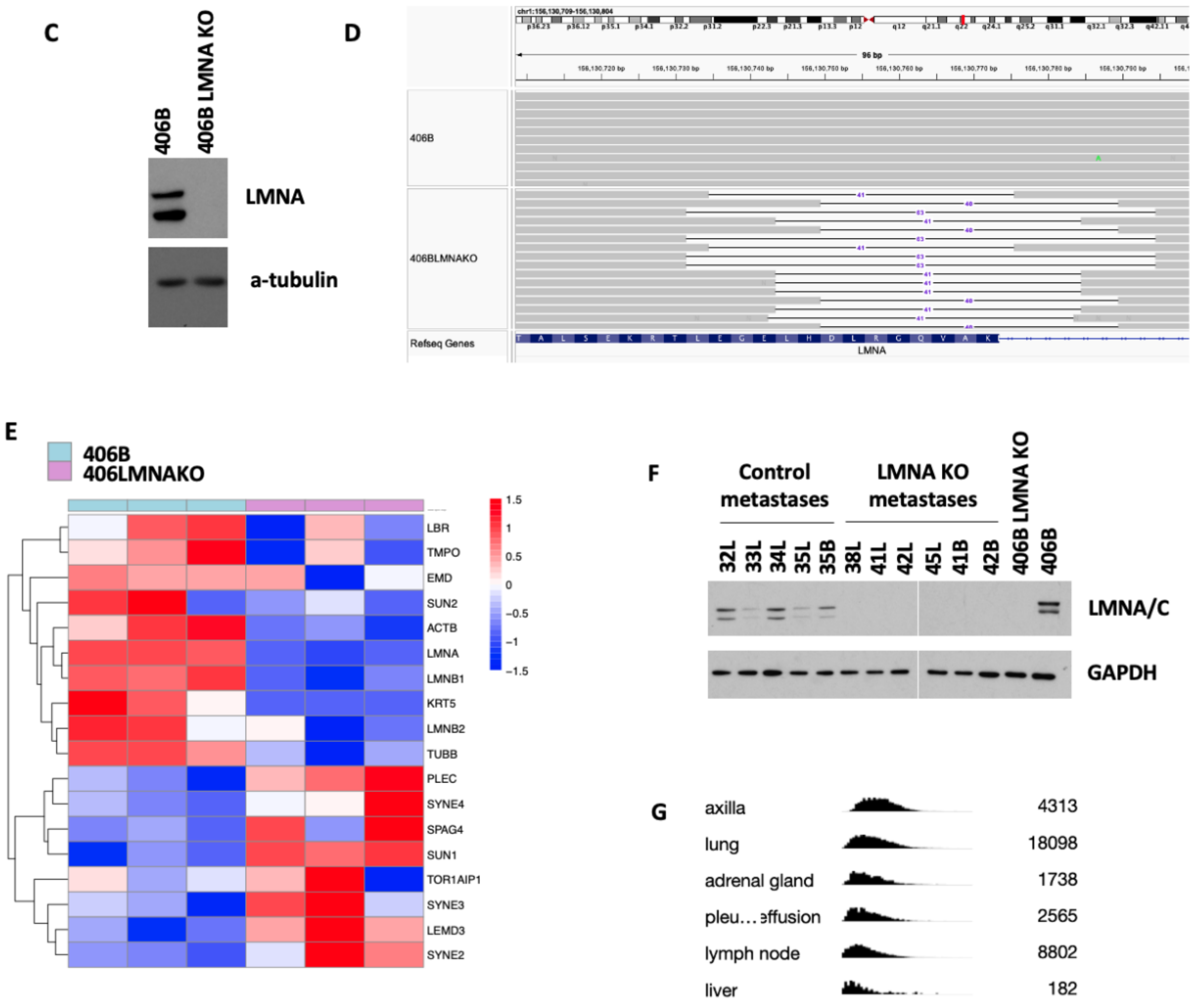

**Figure S5: Dysregulation of cellular structure and LMNA KO in SCLC**

A: H&E illustrating SCLC nuclei in Parent and generation 1 metastases

B: RNA-seq of LINC complex genes in Parent, generation 1, and generation 2 of SCLC metastasis model (shows progressive dysregulation of LINC)

C: Western blot of LMNA KO in 406B

D: Deep amplicon sequencing to identify mutations in 406B LMNAKO

E: Heatmap of gene expression of LINC complex and related genes in 406B and 406B LMNAKO

F: LMNA/C western blot from metastasis cell lines after in vivo LMNAKO metastasis modeling. Samples were run side-by-side on two different blots (separated by white space above) that were developed at the same time.

G: Distribution of LMNA expression of human SCLC metastatic cells [2] expressing, LMNA stratified by site of metastasis

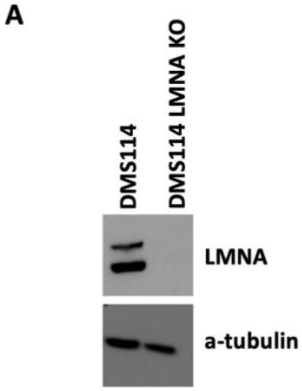

**Figure S6: In vitro examination of LMNA role in nuclear deformability, migration, and metastasis**
A: Western blot of LMNA KO in DMS114

Movies:

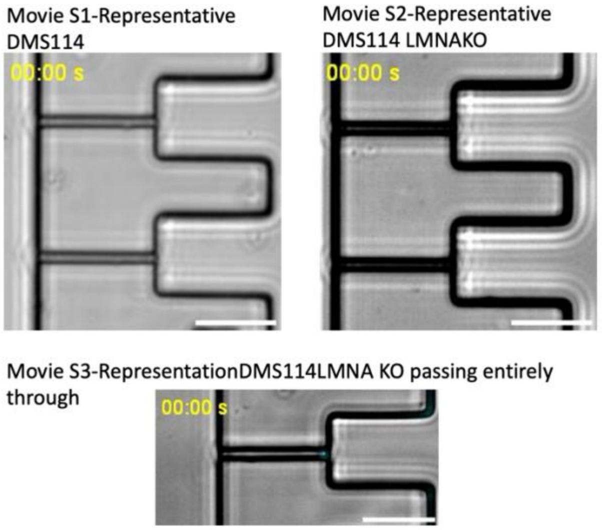

1. Febres-Aldana, C.A., et al., *Rb Tumor Suppressor in Small Cell Lung Cancer: Combined Genomic and IHC Analysis with a Description of a Distinct Rb-Proficient Subset.* Clin Cancer Res, 2022. **28**(21): p. 4702-4713.
2. Chan, J.M., et al., *Signatures of plasticity, metastasis, and immunosuppression in an atlas of human small cell lung cancer.* Cancer Cell, 2021. **39**(11): p. 1479-1496.e18.
